# ZNF768 regulates expression of E2F1 protein to drive G0/G1 transition and cell cycle progression

**DOI:** 10.64898/2026.08.03.742597

**Authors:** Romain Devillers, Benjamin Brisebois, Suparba Roy, Emeline I. J. Lelong, Audrey Poirier, Alona Kolonohuz, Danielle Caron, Christophe Tav, Romain Villot, Frédéric Lessard, Laura Tribouillard, Chantal Garand, Arnaud Droit, Samer Hussein, Philippe Joubert, Mathieu Laplante, Sabine Elowe

## Abstract

Accurate and tightly coordinated cell cycle progression and cell proliferation are critical for development, growth and homeostasis of an organism. Recently, Zinc finger protein 768 (ZNF768) was identified as a transcription factor driving cellular proliferation, in both a p53-dependent and independent manner. ZNF768 interacts with and represses p53 functions to limit cell cycle delay. Independently, ZNF768 promotes the transcription of key regulators of the cell cycle machinery, although the mechanisms through which this occurs remain unknown. Here, we report that ZNF768 protein levels are tightly regulated during the cell cycle, and its depletion leads to cell cycle exit and induction of quiescence. We found that ZNF768 modulates the cell cycle, at least in part, by controlling expression of the major pro-proliferative transcription factor *E2F1* independently of p53 activation. Consequently, depletion of ZNF768, which also represses expression of the key mitotic transcription factor and E2F1 target *FOXM1,* leads to numerous mitotic errors. Supporting these findings, cancer genomics analyses reveal that *ZNF768* expression levels are positively associated with *E2F1* and *FOXM1* expression levels in human tumors, suggesting that cancer cells might use ZNF768 to override cell cycle arrest, sustain proliferation, and promote cancer progression. Altogether, our results reveal that ZNF768 modulates cell cycle entry and proliferation, at least in part by regulating *E2F1* expression.

## Introduction

The regulation of the cell cycle is orchestrated by a network of highly interconnected cyclin-dependent kinases (CDKs) and their cyclin co-factors, as well as rhythmically expressed transcription factors that coordinate orderly progression through proliferation, quiescence, and senescence^1^. One of the best-studied families of transcription factors driving the cell cycle are the E2Fs. These proteins form a core transcriptional circuit crucial for coordinating the oscillatory nature of the cell cycle. Collectively, the E2Fs dictate the timing and fidelity of genome replication and ensure that genetic material is accurately passed through each cell division cycle^2–5^. Recent evidence also highlights the emerging role of C2H2-type zinc finger proteins (ZNFs) in fine-tuning gene expression programs that govern cell cycle progression, proliferation, and senescence^1,6–9^. Notably, many evolutionarily recent ZNF transcription factors display synchronized cell cycle expression and have been shown to actively regulate cell cycle events in human cells, supporting the idea that these factors evolved to provide a regulatory plasticity for cell cycle adaptation^1^. Some notable examples are the primate-specific ZNF519 and simian-specific ZNF274. ZNF519 was found to be highly enriched at promoters of G1/S-regulatory genes and its loss resulted in a decrease in cell cycle progression and cellular proliferation^1^. ZNF274, whose expression peaks in S-phase, has been shown to set replication timing at many of its targets^1^.

Zinc Finger Protein 768 (ZNF768) is a C2H2-type zinc finger transcription factor that evolved in mammals and was recently linked to the control of cell cycle and proliferation^7,9,10^. Its C-terminus is composed of 10 consecutive zinc-fingers with >96% sequence identity in placentals and marsupials. Placental mammals additionally evolved an N-terminus composed of 10-20 heptapeptide repeats reminiscent of those found in the C-terminal domain of the large subunit (RBP1) of RNA polymerase II^7^. ZNF768 preferentially binds to mammalian-wide interspersed repeat (MIR) sequences, retrotransposed DNA elements primarily associated with euchromatic regions and promoters^7^. Mass spectrometry data indicated that ZNF768 interacts with the Elongator complex (Elp1/2/3), implicating this protein in the regulation of transcription elongation through the recruitment of key components of the transcriptional machinery. Functionally, ZNF768 loss was found to impair cell proliferation^7,9^. Indeed, although this transcription factor appeared to control gene expression in a cell-type-specific manner, ZNF768 also regulated the expression of a common core group of genes that support proliferation^7,9^. Furthermore, our recent work demonstrated that oncogenic RAS activation and DNA damage trigger rapid ZNF768 degradation, leading to cell cycle exit and senescence^9^. In line with these observations, ZNF768 overexpression was sufficient to bypass RAS-induced senescence and suppress p53 activation. Supporting a functional link between ZNF768 and p53, ZNF768 was found to interact with and represses p53 activity^9^. Loss of expression of pro-proliferative cell cycle genes, however, preceded the induction of p53 targets, suggesting that ZNF768 might act through both p53-dependent and p53-independent mechanisms to regulate cell proliferation. Altogether, these results support the idea that genotoxic stress downregulates ZNF768 levels to repress the expression of key cell cycle effectors, amplify p53 activation, and reduce cellular proliferation and survival^10^.

Immunohistochemical analyses showed that ZNF768 protein levels are elevated in many human tumors^8^. Tissue microarray performed in a large cohort of lung adenocarcinomas (LUAD) also showed positive associations between ZNF768 and proliferative features including the mitotic score and Ki-67 expression^8^. Supporting the relationship between ZNF768 and cell cycle regulation, ZNF768 knockdown in LUAD cell lines significantly reduced proliferation and downregulated key cell cycle genes^8^. Additionally, patient data reveal frequent ZNF768 amplification or post-transcriptional overexpression, supporting its role as a potential regulator of tumor proliferation^8,10^. In murine models, loss of ZNF768 produced viable animals but resulted in growth delays, embryonic fibroblast proliferation defects, p53 activation, premature senescence, and increased sensitivity to genotoxic stress^11^. Moreover, ZNF768 loss attenuated KRAS^G12D^-driven lung tumorigenesis, underscoring its pro-proliferative and p53-antagonistic role *in vivo*^11^. These studies collectively indicate that loss of ZNF768 reduces proliferation, whereas elevated levels of ZNF768 may be harnessed to promote tumorigenesis in various cancer types.

Although ZNF768 promotes cell cycle progression, the mechanism(s) through which this occurs remains unknown. Here, we show that ZNF768 is dynamically regulated during the cell cycle and is rapidly degraded upon mitotic entry before re-expression in the next G1. ZNF768 regulates the cell cycle by controlling the expression of the key cell cycle transcriptional nodes, E2F1 and consequently its downstream target FOXM1, a major driver of mitotic gene expression. Loss of ZNF768 appears to decrease expression of E2F1 independently of p53 activation, eventually leading to mitotic catastrophes and cell cycle exit. Cancer genomics analyses further demonstrate a positive *in vivo* correlation between ZNF768 expression and E2F1 and FOXM1 levels, supporting the notion that cancer cells may hijack ZNF768 to evade cell cycle arrest, sustain uncontrolled proliferation, and promote malignant transformation. Altogether, these results indicate that ZNF768 functions as a switch modulating cell cycle entry and exit through the regulation of expression of pro- proliferative E2Fs.

## Results

### ZNF768 is a cell cycle-regulated protein that is repressed during mitosis

We and others have previously reported that ZNF768 promotes expression of cell cycle genes and impacts the fidelity of cell division during mitosis; however, the precise mechanisms by which ZNF768 acts in cell cycle regulation remain unclear^7,9^. We first examined ZNF768 protein levels throughout the cell cycle in HeLa cells, a model commonly used in cell cycle studies. Cells were synchronized in G1 using thymidine before being released for up to 10 hours or were enriched in mitosis using the microtubule poisons nocodazole and Taxol. Using these approaches, we observed that ZNF768 protein levels were strongly reduced at mitotic entry, coincident with cyclin B expression (Fig.1A). In agreement, when synchronized in prometaphase using nocodazole, cells exhibited low levels of ZNF768, which then steadily increased after release from the mitotic block, reaching levels comparable to asynchronous cells in the subsequent G1 phase and coinciding with loss of cyclin B (Fig.1B, see also Fig.S1A). To confirm that the loss of ZNF768 protein levels during mitosis is conserved across cell lines, we performed Western blot analyses in both cancerous (HeLa, MDA-MB- 231 and HCT116) and non-cancerous (hTERT-immortalized RPE1, hereafter RPE1) cell lines, either synchronized in mitosis with nocodazole or left untreated. Across different cell lines with diverse genetic backgrounds, ZNF768 protein levels were consistently reduced in mitotic extracts (Fig.1C). In contrast, expression of Transcriptional Repressor CTCF (CTCF), a transcription factor of the same C2H2 zinc finger family as ZNF768, was maintained in mitotic cells. As previously reported, CTCF band migrated slower in the gel in this experiment, suggesting its hyperphosphorylation in mitosis ^12^. These data suggest that loss of ZNF768 protein in mitosis is a common feature of the cell cycle across human cell lines of various genetic backgrounds.

**Figure 1:**
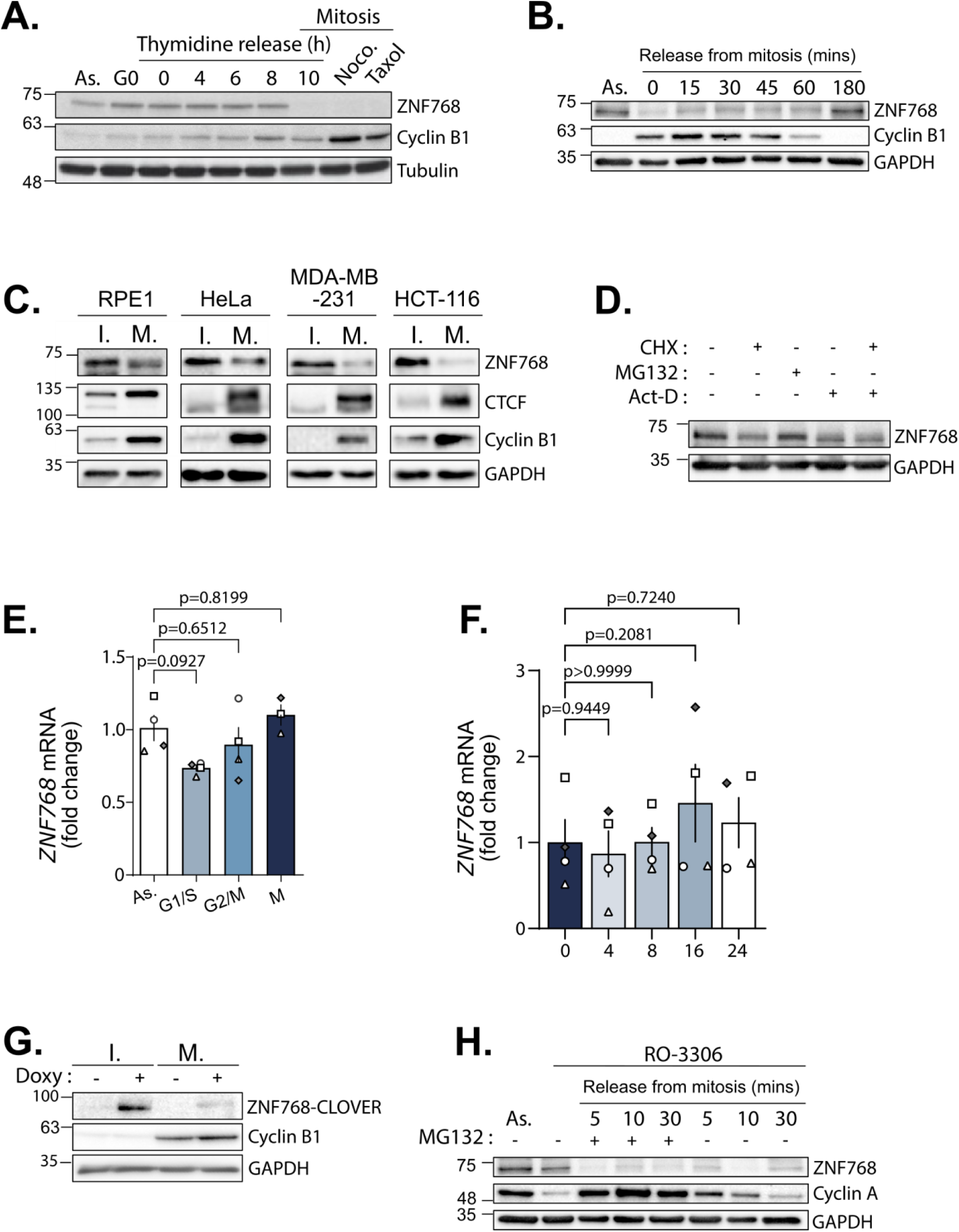
ZNF768 is tightly regulated during the cell cycle and is specifically degraded during mitosis. **A** Cell cycle profile of ZNF768 in HeLa cells that were asynchronous (As.), treated with nocodazole or Taxol, serum starved (G0) or released from thymidine mediated G1 arrest for the indicated timepoints (*n*=1). **B** Asynchronous HeLa cells or cells synchronized in prometaphase using nocodazole, were released and harvested for the indicated time points and lysates were blotted with the indicated antibodies (*n*=3). **C** The indicated cell lines were synchronized in mitosis (M.) using nocodazole or left in interphase (I.) before lysis and Western blotting as indicated (*n*=3). **D** RPE1 cells were treated with 10µg/ml of CHX (cycloheximide), 20 µM of MG-132, 8 µM of act-D (actinomycin D) or a combination for 5hr before lysis and Western blotting as indicated (*n=*3). **E** Relative changes in *ZNF768* mRNA expression in RPE-1 cells synchronized at different stages of the cell cycle, normalized with *B2M* and *GAPDH* (*n=3* for M and *n=4* for others). **F** Relative changes in *ZNF768* mRNA expression in RPE1 cells synchronized in mitosis before release for the indicated time points, (*n=*4). **G** Lysates from HeLa overexpressing doxycycline-inducible mClover-ZNF768 were synchronized in mitosis (M.) with nocodazole or left in interphase (I.) were blotted as indicated (*n = 2*). **H** RPE1 cells were synchronized in G2/M using RO-3306, released in presence of MG-132 and harvested at the indicated time points. Western blots were performed as indicated (*n*=2). In all panels, data represent the mean ± SEM. In panel **E**, **F**, significance was determined by one-way ANOVA followed by Dunnett’s multiple comparisons test.

We next sought to determine the mechanisms regulating ZNF768 levels across the cell cycle. Treatment of asynchronous growing cells with actinomycin D or cycloheximide, two compounds that inhibit transcription and translation, respectively, or with the proteasomal inhibitor MG132, identified only minor changes in ZNF768 protein levels, as revealed by Western blotting (Fig.1D). Moreover, measurement of *ZNF768* expression at different stages of the cell cycle (Fig.1E) or in cells released from mitosis into the subsequent cycle (Fig.1F) showed no significant changes in *ZNF768* mRNA expression, in agreement with the idea that in unperturbed, normally proliferating cells, *ZNF768* expression levels remain relatively constant.

Our previous observations demonstrating rapid proteasomal degradation of ZNF768 in response to genomic insults led us to postulate that the loss of ZNF768 during mitosis might occur at the protein level^9^. In agreement with this, exogenous expression of Clover-tagged ZNF768 revealed that ZNF768 protein levels remained low during mitosis despite increased induction of exogenous ZNF768 in interphase cells (Fig.1G). Moreover, we occasionally observed ZNF768 as a faint doublet band in extracts from mitotic cells, suggesting mitotic-specific post-translational modification (Fig. S1B). To establish whether ZNF768 is a target of proteasomal degradation during mitosis, we synchronized cells at G2/M transition using the CDK1 inhibitor RO-3306, released them in the presence or absence of the proteasome inhibitor MG132, and monitored the expression of ZNF768. Cyclin A, normally subject to degradation upon mitotic entry, was used as a positive control^13^. ZNF768 protein levels showed a small but consistent rescue in the presence of MG132(Fig. 1H).

Overall, our results are in line with the idea that ZNF768 protein levels are tightly regulated across the cell cycle and that ZNF768 is consistently and rapidly lost prior to cell division.

### ZNF768 depletion induces an accumulation of cells in G0/G1

We next sought to characterize the role of ZNF768 during the cell cycle in normal, non-cancerous cells. To this end, we generated RPE1 cell lines expressing previously validated doxycycline-inducible short hairpin RNA (shRNA) targeting two different regions of ZNF768 (sh_ZNF768_1 and sh_ZNF768_2, respectively)^9^. After 48 hours of doxycycline, ZNF768 mRNA and protein levels were both strongly reduced in these cell lines (Fig.2A, 2B). As previously described, we found that loss of ZNF768 impaired proliferation, resulting in an approximately 40-50% decrease in cell number detected as early as 48 hours after induction of ZNF768 depletion, and this attenuated proliferation continued up to 12 days (Fig. 2C). The reduction in cell proliferation was accompanied by the appearance of senescence-like characteristics, including enlarged and flattened cells starting at 4-6 days after shRNA induction. Moreover, ZNF768 was accompanied by a corresponding increase in SA-β-galactosidase activity (Fig. 2D, S2A). Importantly, ZNF768 depletion for 48 hours did not visibly induce SA- β-galactosidase activity indicating early stages of cell cycle exit at this point.

**Figure 2:**
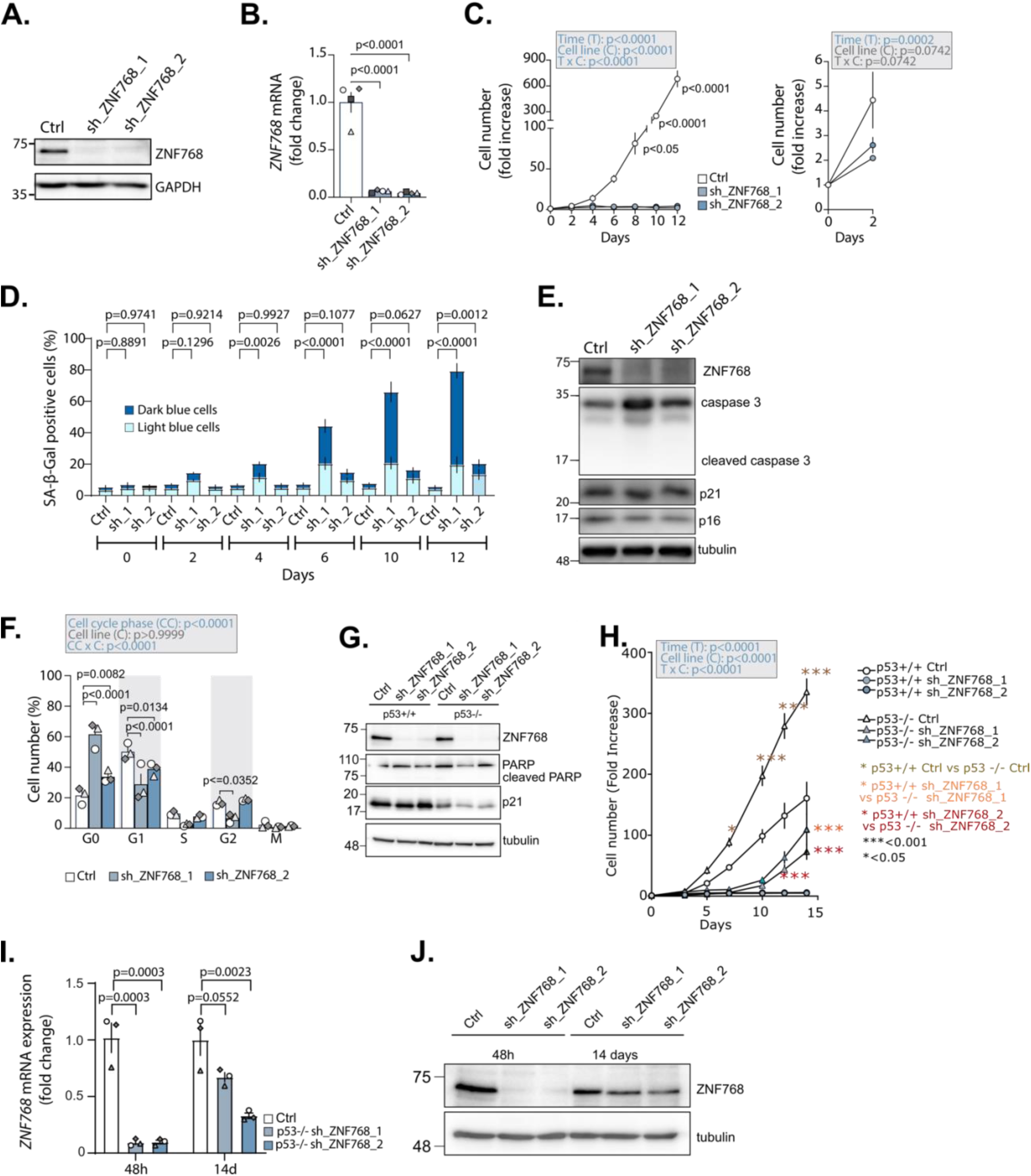
ZNF768 depletion induces an accumulation of cells in G0/G1. Stable, doxycycline inducible RPE1 cells for ZNF768 shRNA knockdown were generated. **A** Western blots and **B** RT-qPCR were performed after 48h induction (*n*=4). **C** Inducible RPE1 cells were counted after ZNF768 knockdown induction for the indicated timepoints. The graph on the right shows growth rate after 2 days of shRNA induction (*n*=3). **D** RPE1 cells were treated as described in (**C**) and SA-β-gal staining was performed. The % of SA-β-gal positive cells is presented (*n*=3). **E** Lysates from cells depleted for ZNF768 for 2 days were blotted with the indicated antibodies (*n*=3). **F** Cell cycle distribution of ZNF768-depleted cells (48h), stained with propidium iodide and FITC anti-Ki-67, measured by FACS (*n*=3). **G** Western blotting of lysates after shRNA mediated depletion of ZNF768 in p53 +/+ and p53-/- RPE1 cells (*n*=3). **H** Cell proliferation after ZNF768 depletion in p53+/+ and p53-/- RPE1 cells (*n*=3). **I** RT-qPCR of *ZNF768* mRNA levels after 48h and 14 days of shRNA induction in matched samples, normalized with *B2M* and *GAPDH* (*n*=3)**. J** Western blotting of ZNF768 protein levels after 48h and 14 days of shRNA induction in p53-/- RPE1 cells (*n*=3). In all quantifications, data represent the mean ± SEM. In panel **B** and **I**, significance was determined by one-way ANOVA followed by Dunnett’s multiple comparisons test. In panel **D**, significance was determined by two-way ANOVA followed by Dunnett’s multiple comparisons test. In panel **F** and **H**, significance was determined by two-way ANOVA followed by Tukey’s multiple comparisons test. In panel **C**, both shRNAs showed similar statistical significance compared with control, for clarity, only one p- value is shown.

Furthermore, at this time point, cells depleted of ZNF768 were not undergoing apoptosis, as evidenced by the absence of caspase cleavage, and exhibited no increase in either p21 (a p53 target gene) or p16 levels, suggesting that the growth arrest was independent of these senescence-inducing factors (Fig.2E). To corroborate the observations of SA-β-galactosidase assay, we used FACS analyses to quantify the proliferative marker Ki-67 at 2 and 7 days following ZNF768 knockdown. After 2 days, approximately 70% of control and sh_ZNF768_2 cells 1 were Ki-67 positive compared with only 30% of cells expressing sh_ZNF768_1. At 7days post-induction, around 80% of control cells remained Ki-67 positive, but this proportion dropped to 40% in response to either sh_ZNF768_1 and sh_ZNF768_2 induction (Fig.S2B). We next examined the cell cycle profile after a 48-hour ZNF768 knockdown using a double-staining approach with propidium iodide (PI) and Ki-67 to distinguish between cell cycle stages^14^. The Ki- 67 antigen, which is rarely detected in G0 phase but highly expressed in proliferating cells, in combination with PI, enables the discrimination of quiescent cells (G0) from proliferative cells and the identification of their cell cycle phases. We found that whereas only 20% of cells accumulated in G0 in the control condition, approximately 60% of sh_ZNF768_1-expressing and 35% of sh_ZNF768_2-expressing cells arrested in G0 after 48 hours of ZNF768 depletion (Fig.2F). Given our previous observations that ZNF768 may function as an inhibitor of p53 signaling, we also explored the effect of ZNF768 depletion on proliferation in RPE1 cells where *TP53* gene (thereafter p53) was inactivated by CRISPR/Cas9^15^. ZNF768 was efficiently depleted in p53-/- cells (Fig.2G). Proliferation was severely impeded up to 10 days after shRNA induction in the p53-/- background, much like depletion in p53 +/+ RPE1 cells, but an uptick in proliferation was observed thereafter (Fig.2H). This renewed capacity for proliferation suggests adaptation despite continual use of fresh doxycycline and shRNA induction. To explore this idea, we induced depletion for 48h and 14 days and compared both *ZNF768* expression and protein levels. We found that while *ZNF768* mRNA and protein expression were markedly reduced at 48h, this reduction was less penetrant at 14 days in the p53-/- background, suggesting that loss of p53 permitted ZNF768 re-expression (Fig.2I, J). In agreement with this, basal levels of ZNF768 expression are higher in p53-/- RPE1 compared to p53+/+ cells (Fig. S2C, D). Overall, these observations confirm our previous reports that ZNF768 depletion results in attenuation of cell cycle progression. These results also suggest a time-window during which the direct effect of ZNF768 loss on cell cycle progression may be explored before the onset of senescence and feedback regulation by p53.

### ZNF768 depletion impairs G0/G1 transition

The observation that ZNF768 depletion led to rapid loss of proliferation, together with our finding that ZNF768 was quickly re-expressed upon cell cycle re-entry after mitosis suggest that ZNF768 may play a role in the G0/G1 transition. To test this hypothesis, we sought to induce a deep quiescent state using serum starvation and assess the ability of control cells and ZNF768-depleted cells to resume proliferation following serum exposure. Recent evidence suggests that quiescence is not a single, uniform state but rather a continuum during which cells gradually enter deeper quiescence, becoming less responsive to growth signals^16,17^. To mimic this, p53+/+ RPE1 cells were cultured for 7 days without serum, during which they retained a fusiform phenotype similar to that of serum-cultured cells, indicating that they had not yet become senescent; although less responsive to growth factors, these cells were still able to exit quiescence and resume proliferation upon serum re-addition (Fig.3A)^16,17^. To examine the role of ZNF768 in exit from quiescence using this approach, p53 +/+ RPE1 lines were starved of serum for 7 days with ZNF768 depletion induced in the 48 hours before serum reintroduction. Cells were subsequently counted every 24 hours for an additional 4 days to assess proliferation (Fig.3B). We found that whereas control cells increased 6-fold in number over 4 days following serum reintroduction, loss of ZNF768 expression prevented cells from re-entering the cell cycle and further proliferating (Fig.3C). To explore this growth arrest at the molecular level, we performed Western blot analysis of cell extracts collected at different time points after serum starvation and re-entry. As expected, in all cell populations, serum starvation decreased protein expression of key cell cycle and proliferative markers in all cell lines (e.g. Ki-67, pRB, E2F1; see prolif. vs. D8). Following serum reintroduction, levels of proliferation markers increased in control cells to levels comparable or superior to proliferating cells, whereas in the absence of ZNF768, proliferation marker levels remained low (sh_ZNF768_2) or virtually undetectable (sh_ZNF768_1) even after reintroduction of serum (Fig.3D). As shown previously, no striking differences in p53 or p21 protein levels were observed upon ZNF768 depletion under these conditions (Fig. 3D). In contrast to our observations in p53+/+ cells, loss of p53-/- permitted proliferation in ZNF768 depleted cells, albeit at a lower rate, presumably as a consequence of ZNF768 re-expression (Fig. 3E, 2I, J). Collectively, these results indicate that immortalized, non-carcinogenic RPE1 cells require ZNF768 to re-enter the cell cycle and proliferate after prolonged arrest (7 days), but that in the absence of p53, proliferation can eventually be reinitiated in ZNF768-depleted cells.

**Figure 3:**
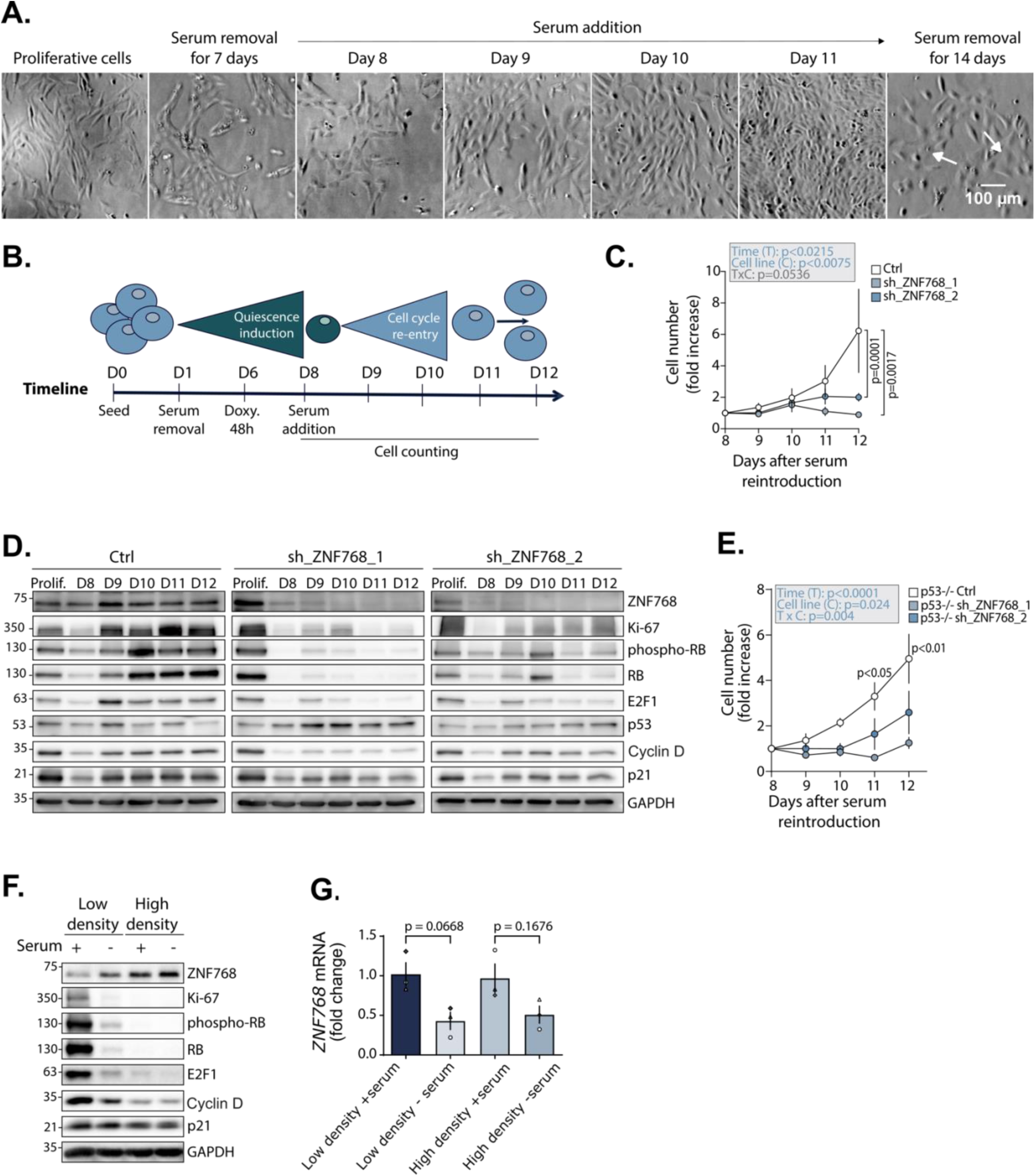
ZNF768 depletion impairs G0/G1 transition. **A** Morphology of cells cultured in different serum conditions visualized by phase contrast microscopy. White arrows highlight cells exhibiting a more flattened and enlarged morphology. **B** Schematic representation of experiments performed in panel (**C-E**). **C** Inducible p53+/+ RPE1 ZNF768 shRNA cells were cultured as in **B** without serum for 7 days then exposed again to serum after a 48 h ZNF768 knockdown and counted (*n*=3). **D** Inducible RPE1 cells were treated as described in **B, C** and Western blots were performed (*n*=3); Proliferation conditions (*Prolif.*) correspond to corresponding cell lines without knockdown induction. **E** Inducible p53/- RPE1 ZNF768 shRNA cells were treated as in **C** and counted (*n*=3). **F** RPE1 cells were cultured at low or high density and with or without serum for 7 days. Western blots were performed as indicated (*n*=3). **G** RPE1 cells were cultured as described in **F** and RT-qPCR were performed to measure *ZNF768* mRNA levels (*n*=3). In all panels, data represent the mean ± SEM. In panel **C** and **E**, significance was determined by two-way ANOVA followed by Tukey’s multiple comparisons test. In panel **G**, significance was determined by two-way ANOVA followed by Tukey’s multiple comparisons test.

As presented in Figure 1, we observed a sharp decrease in ZNF768 at the “end” of the cell cycle during mitosis. This, together with observations that ZNF768 is rapidly degraded upon cellular insults such as DNA damage or oncogenic stress^9^, suggests that ZNF768 loss might be a prerequisite to efficiently drive cell cycle exit. To test this idea, we sought to understand whether ZNF768 expression can be modulated by forced switching between proliferating and non-proliferating state. For this, RPE1 cells were cultured with or without serum for 7 days and, in both conditions, at low and high density to mimic proliferative and growth-arrested state, respectively. At low confluency, proliferation markers such as Ki-67, Cyclin D1, E2F1 and pRb were low in the absence of serum, but their expression increased in the presence of serum, as anticipated. Surprisingly, ZNF768 protein levels showed the opposite pattern, being low in the presence of serum and increased under conditions that induced quiescence (low density without serum, and high density with or without serum), consistent with the idea that cell cycle exit may trigger relatively high ZNF768 expression preemptively as a mechanism to facilitate subsequent cell cycle re-entry (Fig.3F). Moreover, we found that mRNA levels behaved differently than protein levels and were consistently higher in the presence of serum regardless of cell density (Fig.3G). Although these observations seem contradictory, they support the idea that ZNF768 protein levels are tightly controlled across the cell cycle.

### ZNF768 regulates key cell cycle genes

ZNF768 was identified as a transcription factor that binds mammalian-wide MIR sequences and promoter regions to control gene expression^7^. To understand how ZNF768 modulates proliferation and cellular quiescence/senescence, we performed RNA-seq to characterize the gene expression profile following 72 hours of ZNF768 depletion in p53+/+ hTERT-RPE1 cells. This time point was chosen to capture a signature induced by early effects of ZNF768 loss prior to a deepening quiescence and onset of senescence. Analysis of gene expression revealed 1287 and 1556 genes significantly downregulated by sh_ZNF768_1 and sh_ZNF768_2 respectively, whereas 821 and 1098 were found to be upregulated in sh_ZNF768_1 and sh_ZNF768_2, respectively (Fig.4A). Of these, 326 genes were commonly downregulated whereas 179 genes were upregulated after ZNF768 depletion with both shRNAs (Fig 4B). Gene set variation analysis (GSVA) revealed a strong decrease in the expression of cell cycle related genes after ZNF768 depletion with both shRNAs, (Fig.4C). For instance, genes encoding proteins involved in the regulation of cell cycle phase transitions (e.g., *CDK2, CCNE2*), DNA replication (e.g., *MCM2-6, PCNA*), genome stability (e.g. *BARD1, BLM, BRCA1, BRCA2, FANCD2, FANCE, FANCG, EZH2, MYB, MYBL2*) and transcription factors governing the cell cycle (e.g. *E2F1, E2F2, FOXM1*) were significantly repressed in both ZNF768 knockdown conditions compared to control, indicating a coordinated attenuation of proliferative transcriptional programs (Fig.4A, D, Supp. table 1). The observation that cell cycle genes were the most prominently downregulated targets of ZNF768 was further supported by gene ontology analysis^18^. In contrast, genes that were upregulated following ZNF768 loss showed substantially weaker GO term enrichment (Fig.S3A, B, Supp table 2,3). In strong agreement with our p53-independent cell cycle regulation, we did not observe a marked induction of canonical p53 target genes, nor upregulation of the senescence-associated secretory phenotype (SASP) genes (Fig.4A, C). To complement this analysis, we used the gene expression profile upon ZNF768 overexpression in HEK293T cell line using iLincs, a publicly available resource providing the expression profile of almost 1000 genes (L1000 assay) in response to various perturbagens^19^. Remarkably, this analysis demonstrated that ZNF768 overexpression resulted in an increase in the number of cell cycle regulatory genes including *MYC, CCNE2, PLK1, PCNA, CDK1, CCNB1, AURKB* and *AURKA* (Fig.4E). These results were further confirmed by gene ontology (GO) annotation analysis which demonstrated that “regulation of cell cycle processes” as the most significantly enriched term (Fig.4F). Overall, these observations agree with previous results and indicate that ZNF768 expression is positively associated with the expression of genes involved in proliferation and cell cycle regulation.

**Figure 4:**
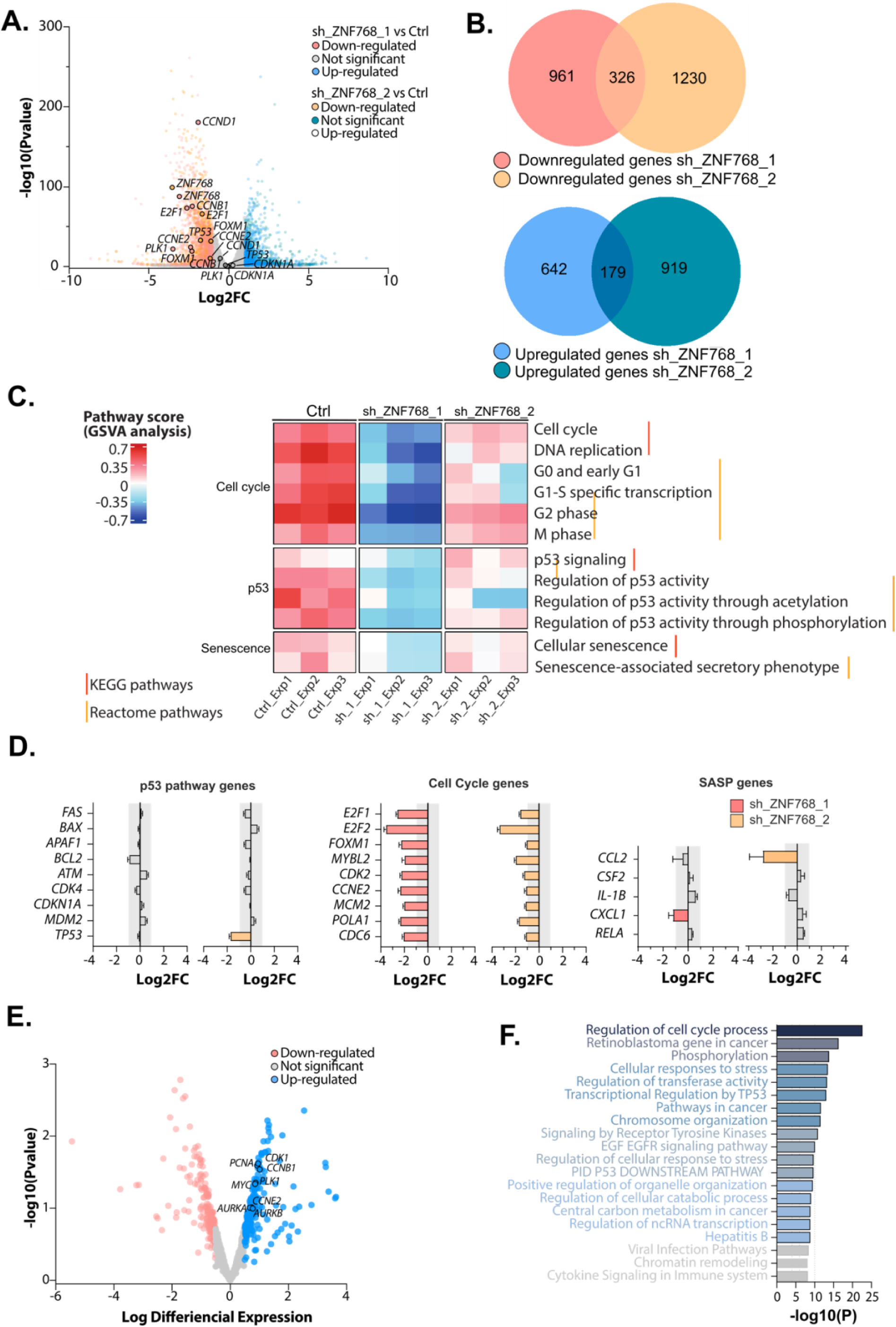
ZNF768 regulates key cell cycle genes. **A** Volcano plot of RNA-seq analyses after ZNF768 depletion in p53+/+ RPE1 cells (*n*=3). RNA was isolated 72 h post induction. Colored dots represent genes significantly (*p*-value <0.05) up or down regulated with a Log2FC <-1 or >1. **B** Venn diagram of downregulated and upregulated genes after depletion of ZNF768 with sh_ZNF768_1 and sh_ZNF768_2. **C** Heatmap representing results of GSVA analysis and enrichment of KEGG and Reactome pathways. **D** Bar graph depicting mRNA expression levels of p53, cell cycle, and SASP related genes in the experiment described in **A**. Colored bars indicate genes that are downregulated (Log2FC < −1). **E** Volcano plot of differentially expressed genes in response to ZNF768 overexpression from the iLincs dataset. The x-axis represents the log differential expression, and the y-axis represents the significance of gene expression in overexpressing versus control cells. Colored dots represent genes up (blue) or down (peach) regulated with a LogDE <-0.5 or >0.5. **F** Gene ontology analysis performed with Metascape of the list of genes upregulated in dataset described in **E**.

### ZNF768 regulates the expression of the E2F family of transcription factors

ZNF768 has been proposed to function as a master regulatory transcription factor positioned upstream of a hierarchical network of other transcription factors ^20,21^. We therefore next sought to determine whether ZNF768 directly regulates the transcription of cell cycle regulatory genes or whether their observed downregulation might be occurring by the intermediary of other transcription factors. Focusing on genes commonly downregulated following ZNF768 depletion with both shRNAs, Transcription Factor Targets and Transcriptional Regulatory Relationships Unraveled by Sentence-based text mining (TRRUST) analysis revealed that genes downregulated in response to ZNF768 knockdown were mainly known targets of the cell cycle regulatory transcription factor E2F1 (Fig.5A, B)^22^. Similarly, TRRUST analysis of the iLincs dataset identified E2F1 as the predominant transcription factor associated with upregulation of genes following ZNF768 overexpression (Fig.5C). Analysis of ChIP-seq data from multiple independent studies for ZNF768 binding in U2OS, Raji, HEPG2 and HEK293T cells all revealed that ZNF768 binds to virtually identical sites at the *E2F1* locus suggesting that E2F1 is a common target of ZNF768 across different lineages (Fig.5D). A recent study identified ZNF768 as an mRNA binding protein^23^. However, E2F1 transcripts were not strongly enriched in the ZNF768 interactome by iCLIP (Fig.S4A) and ZNF768 depletion did not result in changes in E2F1 mRNA stability (Fig.S4B)^24^. In addition to E2F1, ChIP-seq analysis above revealed ZNF768 binding peaks at the pro-proliferative *E2F2* locus, but not other E2Fs family members (Fig.S4C). Taken together, these results suggest a direct regulation of the expression of *E2F1* and potentially *E2F2* by ZNF768.

**Figure 5:**
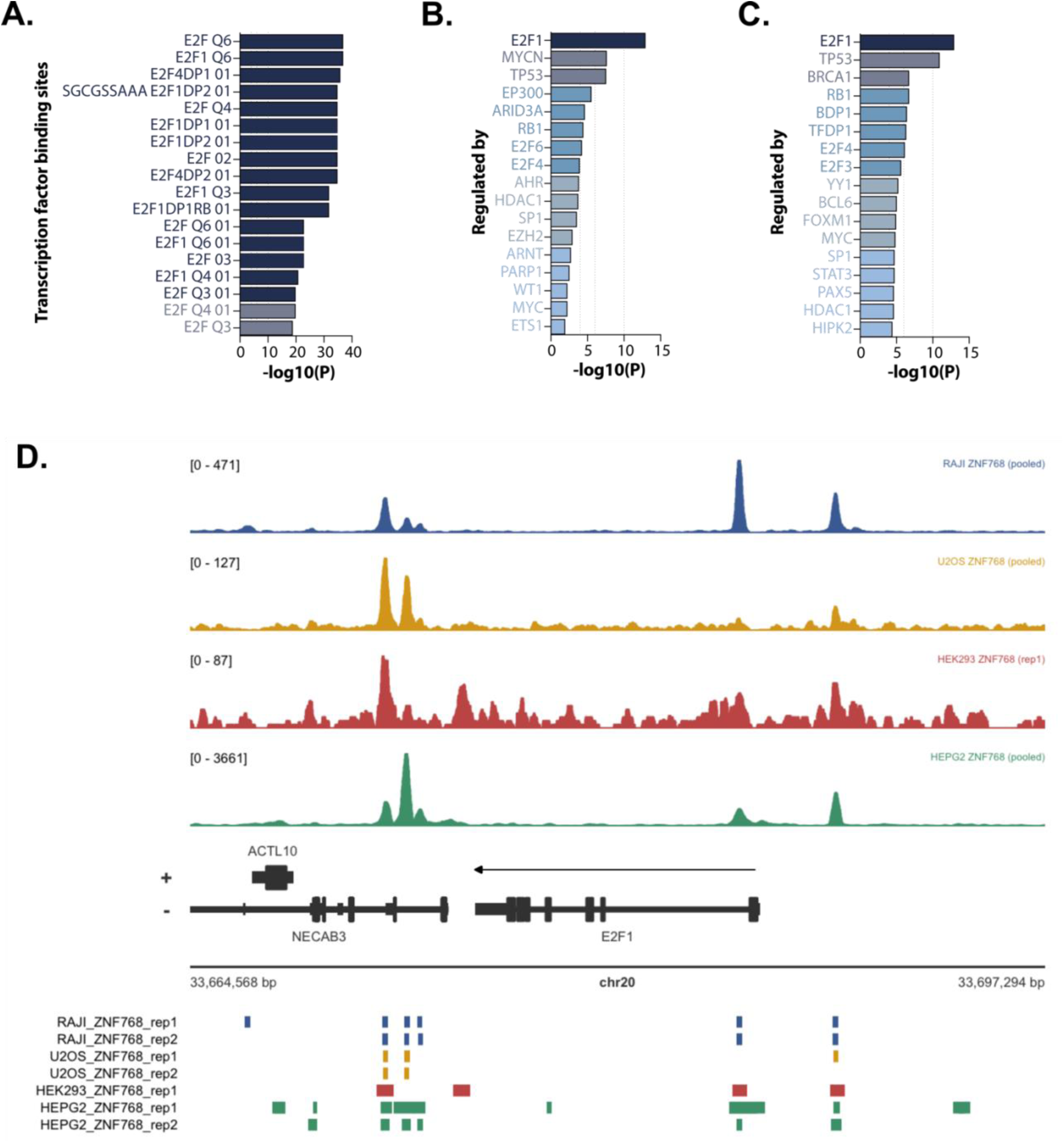
ZNF768 regulates E2Fs pathway. **A** Metascape transcription factor target analysis of genes downregulated in response to ZNF768 depletion. **B** TRRUST analysis of genes identified by RNA-seq that were downregulated in response to ZNF768 depletion. **C** TRRUST of genes upregulated after ZNF768 depletion. **D** ZNF768 binding at the E2F1 locus identified in ChIP-seq data in different cell lines. Identified peaks (by MACS (RAJI, U2OS and HEK293) or SPP (HEPG2)) are shown as rectangles.

### ZNF768 regulates the transcription of key cell cycle genes

E2Fs are recognized as major transcriptional regulators of the cell cycle forming a core transcriptional network driving the expression of many genes involved in cell cycle progression and genome integrity^4^. Deregulation of their transcriptional activity, which has been postulated to be an almost universal event in cancer development^3,4,25^, occurs primarily by perturbation of the CDK-RB-E2F axis^3,4^. E2F1 is the prototypical activator E2F; its expression is induced in mid-late G1, and its activation after release from RB-mediated inhibition is the key event of the restriction checkpoint, G1/S transition and cell cycle commitment^5,16^. Given the critical role of E2F1 in the progression of the cell cycle, and the loss of E2F1 target gene expression in ZNF768- depleted cells, we sought to further investigate the link between these two transcriptional regulators. ZNF768 knockdown in both p53+/+ and p53-/- RPE1 cells revealed reduced protein levels of E2F1 (Fig.6A). In agreement with the results presented above in Fig 2E, we found no activation of p21 upon ZNF768 depletion in the p53+/+ cells, indicating a lack of p53 activation (Fig.S5A). Moreover, pRb, which is subject to feedback regulation by E2F1, also exhibited reduced expression in ZNF768 depleted cells (Fig. S5A)^26^. In p53-/- cells, although ZNF768 depletion demonstrated a strong downregulation of *E2F1* expression after 48 hours of depletion, depletion for 14 days revealed restoration of *E2F1* gene and protein expression despite consistent induction of shRNA, mirroring the restauration of ZNF768 protein expression (Fig. 6B,C and Fig. 2J). These data demonstrate that although ZNF768 loss initially suppresses E2F1 expression independently of p53, cells ultimately adapt and regain *ZNF768* and consequently *E2F1* expression over time in the absence of p53.

**Figure 6:**
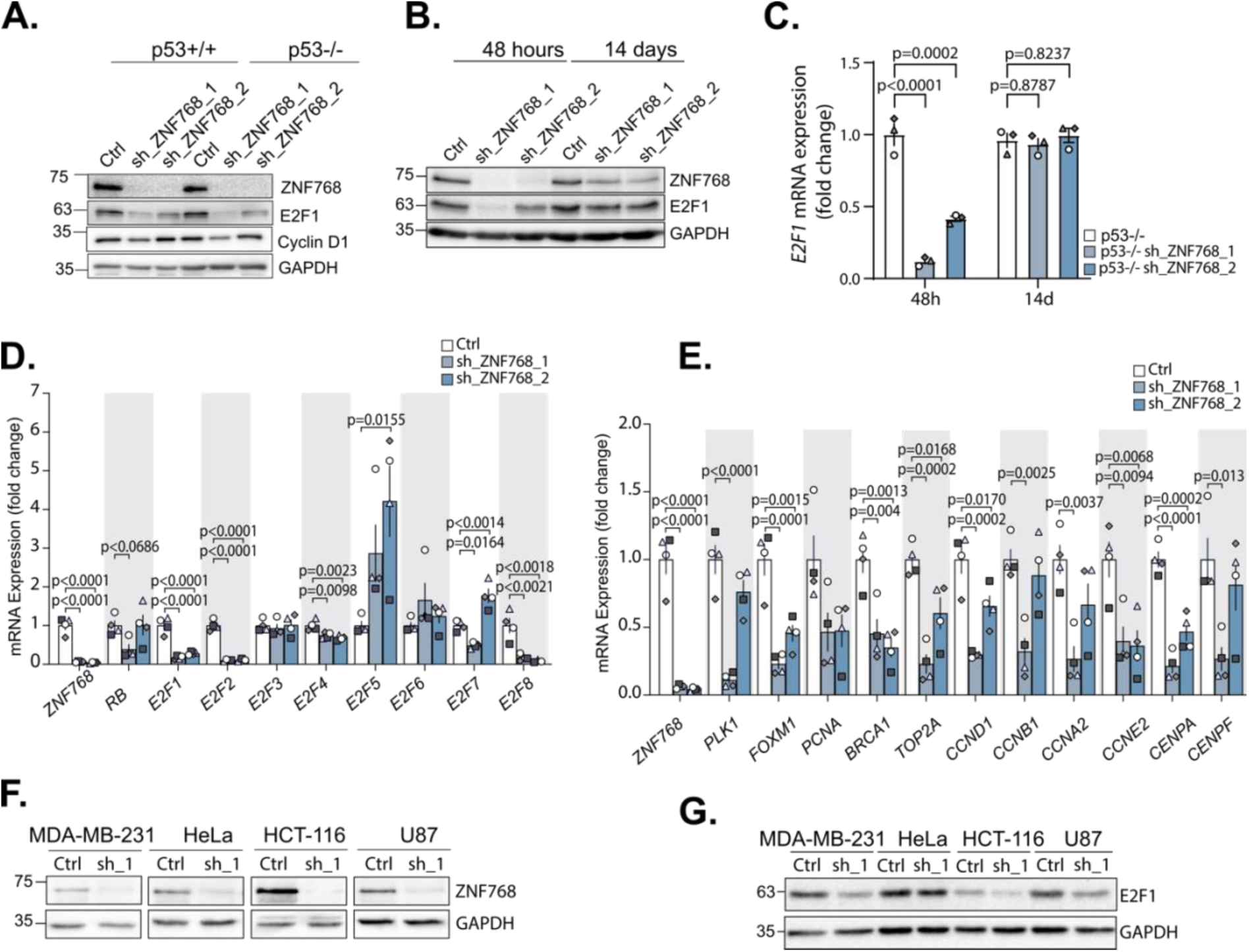
ZNF768 regulates the transcription of key cell cycle genes. **A** RPE1cells were depleted of ZNF768 for 48 hours and blotted for the indicated proteins (*n*=3). **B** p53-/- RPE1 cells were depleted of ZNF768 for 48 h or 14 days and blotted for the indicated proteins (*n*=3). **C** *E2F1* mRNA change in p53-/- cell depleted of ZNF768 after 48 hours and 14 days of depletion, normalized with *B2M* and *GAPDH* (n=3). **D** Expression of E2F family transcription factors in p53+/+ cells after ZNF768 depletion (*n*=4). **E** Expression of E2F target genes in p53+/+ cells after ZNF768 depletion (*n*=4). **F, G** Stable, inducible sh_ZNF768_1 cell lines were generated in MDA-MB-231, HeLa, HCT-116 and U87. Lysates were subjected to Western blotting as indicated after 48 hours of shRNA induction (*n*=3). In all graphs, data represent the mean ± SEM. In panel **C**, **D** and **E**, significance was determined by one-way ANOVA followed by Dunnett’s multiple comparisons test.

We next explored the effect of ZNF768 depletion on the expression of E2F family of transcription factors in general. In agreement with the above ChIP-seq and RNA-seq analysis, in p53+/+ RPE1 cells, expression of *E2F1* and *E2F2* was strongly reduced after ZNF768 depletion with either shRNA, as was expression of the atypical family member *E2F8*, a known E2F1 target gene^27,28^. In contrast, expression of the transcriptional repressor *E2F5* was upregulated (Fig.6D). In support of the loss of pro- proliferative E2F1 expression, we also observed a decrease in the expression of numerous E2F targets in ZNF768-depleted cells including the key cell cycle relevant targets *FOXM1 and PLK1* (Fig 6E). We next generated a panel of cancer cell lines with inducible ZNF768 depletion using sh_ZNF768_1 to determine whether reduced E2F1 expression is a general consequence of ZNF768 loss (Fig.6F). Western blot analysis confirmed a marked decrease in E2F1 protein levels following ZNF768 knockdown in most cell lines (Fig. 6G). This was further supported by RT-qPCR analysis, which demonstrated not only decrease in *E2F1* gene expression but also decreased expression of key E2F target genes, including *FOXM1*, *PLK1*, and *CDK1* across all cell lines (Fig.S5B). These results are consistent with our data in RPE1 cells showing the regulation of E2F1 expression by ZNF768 and suggest that depletion of this zinc finger protein may drive cell cycle exit because of attenuated expression of an E2F1-driven gene expression program.

### Mitotic defects linked to ZNF768 depletion are associated with reduced expression of FOXM1 and PLK1

Aberrant mitosis is a major characteristic of cells depleted of ZNF768 suggesting that downstream transcriptional targets of ZNF768 may be involved in cell division^9^. Cell cycle transcriptional progression is tightly controlled by the sequential activation of two waves of gene expression circuits, starting with RB-E2F at G1/S and followed by the DREAM MYB:MuvB (MMB):FOXM1 factors during G2/M transition; expression of FOXM1 is at least in part dependent on E2F1 activity and functions in G2 and G2/M to promote a transcriptional program that includes key mitotic regulators such as polo-like kinase 1 (PLK1), Cyclin B1, Cyclin dependent kinase 1 (CDK1), Aurora A and Aurora B kinases to ensure the proper execution of mitosis and the fidelity of cell division^29^. In agreement with the loss of *PLK1* and *FOXM1* expression in ZNF768- depleted cells, we found that both PLK1 and FOXM1 protein levels were generally decreased in mitotic extracts from our inducible cell lines after ZNF768 knockdown (Fig.7A and S5B). Moreover, immunofluorescence analyses demonstrated a significant reduction of PLK1 levels at kinetochores, (a principal site of its mitotic activity) across all cell lines examined (Fig.7B, 7C).

**Figure 7:**
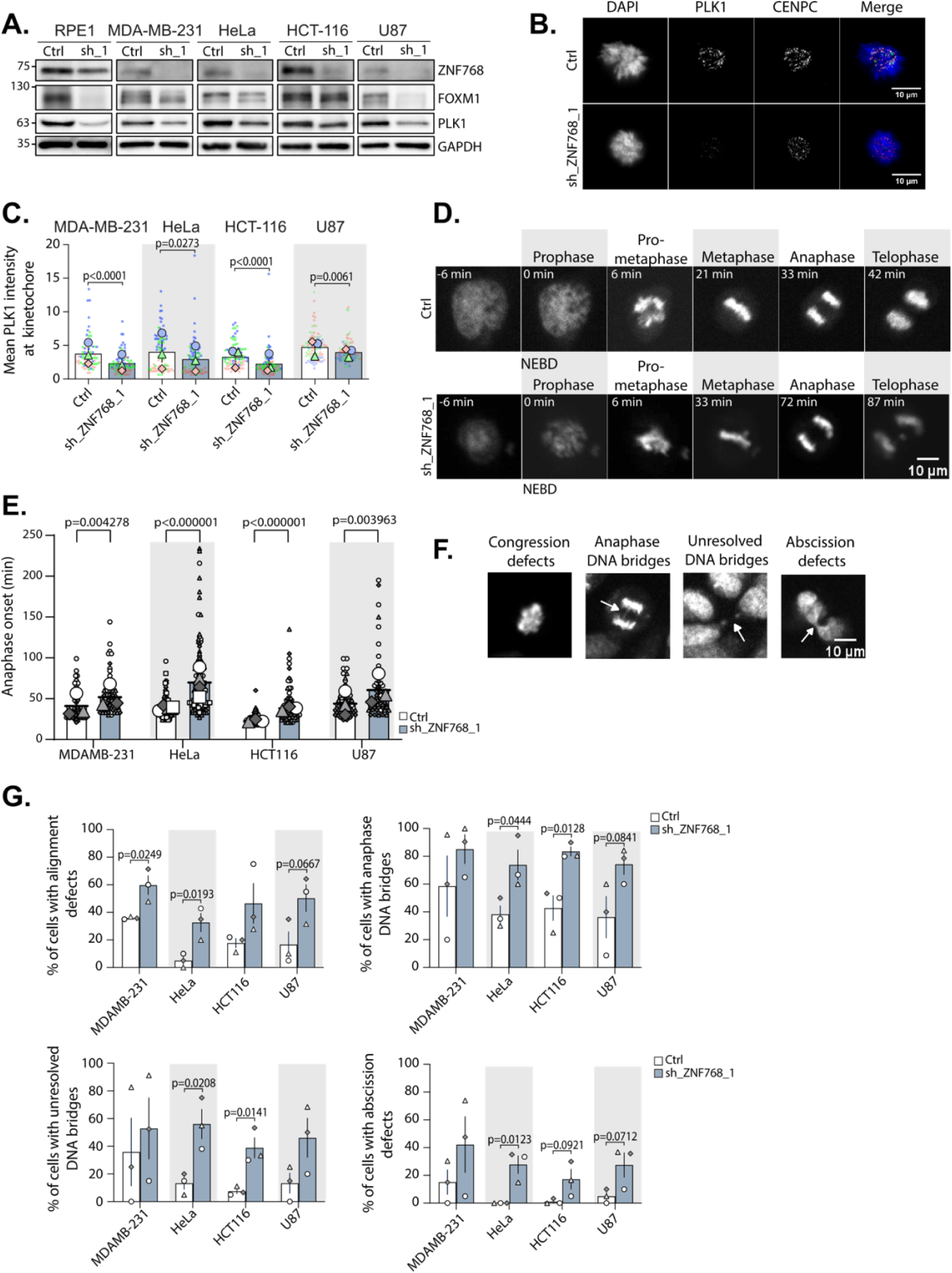
Loss of ZNF768 induces loss of FOXM1, PLK1 and consistent mitotic defects. **A** Western blots were performed on mitotic lysates generated from the panel of cell lines depleted of ZNF768 with inducible sh_ZNF768_1 (*n*=3). **B** Representative immunofluorescence image of mitotic inducible HeLa cells after 48 hours doxycycline induction. Cells were stained for PLK1 (green), CENPC (grey) and with Hoechst (blue). **C** Quantification of the average relative intensity of PLK1 at kinetochores from (**B**) using previous inducible cancer cell lines (*n*=3). **D** Representative stills of time-lapse movies of inducible HeLa cells after a 48 hours ZNF768 knockdown. For imaging, cells were synchronized in mitosis after release from a double thymidine block and incubated with SiR-DNA to visualize the chromatin. **E** Quantification of the average nuclear envelope breakdown to the anaphase onset duration in the indicated cell lines (*n*= n=3 for all cell lines except for the HeLa, n=4). **F** Representative live cell images of mitotic defects observed after 48 hour ZNF768 depletion in HeLa cells. **G** Quantification of mitotic defects listed in **F** in MDA-MB-231, HeLa, HCT-116 and U87 cell lines after 48 hour ZNF768 depletion (*n*=3). In all panels, data represent the mean ± SEM. In panel **C**, **E** and **G**, significance was determined by unpaired *t* test.

Next, we performed live cell imaging in our panel of cell lines after 48 hours of ZNF768 depletion and found that mitotic duration was consistently prolonged in all cell lines examined, consistent with partial PLK1 depletion (Fig.7D, E). Moreover, live cell imaging revealed multiple mitotic defects associated with ZNF768 loss that are consistent with attenuated activity of key mitotic regulators such as PLK1 and Cyclin B-CDK1 ^30–32^. These defects included increased congression defects during chromosome alignment, as well as increased anaphase bridges, abscission defects during cytokinesis, and unresolved DNA bridges in newly born daughter cells in all cell lines examined (Fig.7F, G). Altogether, these results show that ZNF768 loss induces a decrease in key mitotic effectors and FOXM1 target genes such as PLK1, leading to an increase in the number of mitotic defects and thus impaired execution of mitosis.

### *ZNF768* correlates with proliferative gene expression in human cancers

Given the importance of ZNF768 in supporting cell proliferation, we next sought to define whether *ZNF768* expression correlates with previously identified *in vitro* key cell cycle genes in human cancers. Focusing our efforts on lung adenocarcinoma (LUAD), a disease previously associated with ZNF768 overexpression^8,9,11^, we used human transcriptome cancer data from TCGA PanCancer Atlas to explore the correlation of ZNF768 with the major transcriptional drivers of the cell cycle. This work revealed that *ZNF768* expression is significantly correlated with *E2F1, FOXM1,* and *PLK1* expression in LUAD (Fig.8A). These findings were further confirmed using samples collected from our validation cohort of LUAD patients that were divided into low and high ZNF768 groups based on protein and mRNA levels (Fig.8B). Confirming our previous observations in cancer cell lines (Fig.6G, S5B), the high ZNF768 group was characterized by a significant increase in the expression of key cell cycle genes compared to the low ZNF768 group (Fig.8C). Western blot analyses validated the above findings and demonstrated that relatively high expression of ZNF768 correlated with elevated FOXM1, E2F1 and PLK1 protein levels (detectable in 3/3 samples) compared to the low ZNF768 group (detectable in 1/3 samples) (Fig.8D). Finally, Kaplan-Meier survival analysis ^33^ revealed a significantly reduced overall survival in patients with LUAD in high *ZNF768* expression group compared to low *ZNF768* expression group (log-rank test, p < 0.05) (Fig.8E). Collectively, these results are consistent with our *in vitro* data showing the importance of ZNF768 for the expression of several cell cycle genes (and thus proliferation) by increasing *E2F1* expression. These results also highlight the intriguing possibility that elevated ZNF768 expression observed in cancer may contribute to hyperproliferation of cancer cells and thus tumor development.

**Figure 8:**
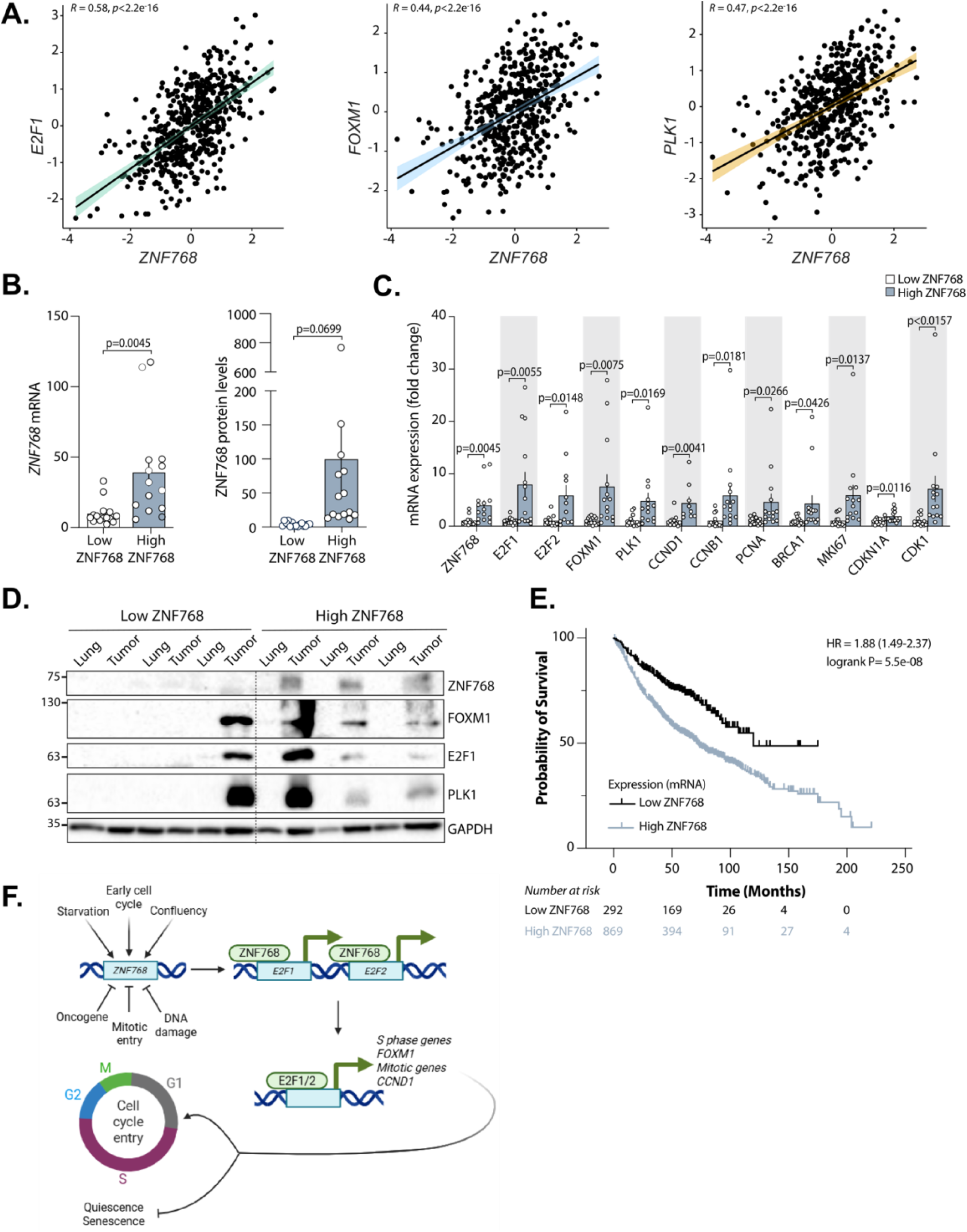
ZNF768 expression correlates with proliferative gene expression in human cancers. **A** Correlation of *ZNF768* and *E2F1, PLK1* and *FOXM1* gene expression analysis from TCGA data bank. **B** Quantification of ZNF768 protein and mRNA levels in low- and high-expressing ZNF768 lung adenocarcinoma sample groups, normalized with *ACTA1*. **C** Gene expression analysis performed by RT-qPCR in sample groups based on ZNF768 levels, normalized with *ACTA1* (*n*=10-14) described in **B**. **D** Western blots performed on samples described in (**C**) (*n*=3). **E** Kaplan–Meier survival curves showing the overall survival of patients diagnosed with lung adenocarcinoma, stratified according to high vs. low expression of *ZNF768*. The survival difference between groups was assessed using the log-rank test. **F** Model of cell cycle regulation by ZNF768. See discussion for details. In all panels, data represent the mean ± SEM. In panel **A**, Pearson correlation analysis was used to determine the strength and direction of associations between variables. Statistical significance was set at p < 0.05. In panel **B** and **C**, significance was determined by unpaired *t* test.

## Discussion

Previous reports have highlighted the importance of ZNF768 in cell cycle progression and proliferation but the exact mechanisms of how this occurs remains unclear. Here, we demonstrate that ZNF768 is a novel regulator of the expression of the master cell cycle transcription factor E2F1. Loss of ZNF768 resulted in reduced expression of the pro-proliferative transcriptional program normally activated downstream of E2F engagement including reduced expression of the E2F1 target FOXM1 (a major mitotic transcriptional regulator) and the FOXM1 target PLK1. The ultimate outcome of this is a rapid decrease in proliferation and cell cycle exit.

Our analysis of ChIP-seq data from different sources and cell lines has highlighted the ability of ZNF768 to bind the *E2F1* promoter. A recent survey of Zinc finger proteins using iCLIP has revealed that many proteins in this group (including ZNF768) bind to RNA and function in diverse post-transcriptional regulatory processes^23^. However, E2F family members, including E2F1 and E2F2, were not appreciably bound by ZNF768 (Fig. S4A). Moreover, we found no significant effect of ZNF768 depletion on E2F1 mRNA stability (Fig.S4B). While binding of ZNF768 in the promoter region of the pro-proliferative *E2F1* and *E2F2* genes across multiple cell lines suggests direct regulation, further studies will be required to demonstrate this definitively.

Cell cycle progression was initially attenuated after ZNF768 depletion in both p53 +/+ and p53-/- cells. Indeed, under conditions of reduced proliferation after ZNF768 depletion, we found no evidence of p53 activation after ZNF768 depletion, including expression of the cell cycle inhibitor p21 (Fig. 2E and S5A). In p53-null cells, an initial downregulation was followed by a compensatory response that enabled cells to resume proliferation, albeit at attenuated levels compared to control. This rebound was associated with an increase in ZNF768 and E2F1 expression despite sustained shRNA induction (Fig. 2J, 6B). The re-emergence of ZNF768 expression may reflect selective pressure favoring its restoration in the presence of growth promoting signals (Fig. 3D). Because ZNF768 can repress p53 activity, ZNF768 and p53 may be involved in a reciprocal regulatory relationship, in which p53 represses the expression of its own inhibitor (Fig S2C, D). This would constitute a double-negative feedback loop and is reminiscent of the feedback between the deacetylase sirtulin-1 (SIRT1) and p53 in which p53 represses expression of negative regulator ^34,35^. Given that ZNF768 acts as a pro-proliferative factor, this circuit may function as a switch-like module governing proliferative capacity at the beginning of each cell cycle. In a cancer context, the reported amplification of ZNF768 could simultaneously enhance proliferative signaling and suppress p53 activity, thereby promoting cell cycle progression^9^. This regulatory dynamic could be further exacerbated in the absence of p53, a condition permissive to sustained upregulation of *ZNF768* mRNA. Importantly, loss of p53 alone was insufficient to elevate ZNF768 protein levels, at least in otherwise non-cancerous RPE1 cells, despite a marked increase in its mRNA expression (Fig.S2C). This implies the presence of compensatory post-transcriptional mechanisms that constrain ZNF768 protein abundance that remain to be explored.

A surprising result was the discordance between ZNF768 and E2F1 protein levels in non-proliferative cells where relatively high levels of ZNF768 were found compared to cells actively undergoing proliferation (Fig.2F). Although counterintuitive, the increase in ZNF768 during contact inhibition or serum starvation suggests that ZNF768 may accumulate to prime cells for G1 re-entry (and potentially G1/S transition) by augmenting E2F activity. Quiescent cells normally have low levels of E2F activators (for example Cyclin D1, Cyclin E, Cdk2) and elevated CDK inhibitors (such as p21 and p27), which repress DNA-replication genes including E2F1 ^36^, ^37,38^. The balance between G1 cyclins and CDK inhibitors such as p21 governs the decision between continued cell cycle progression and entry into quiescence ^39,40^. By promoting E2F1/2 expression, relatively elevated levels of ZNF768 in quiescence could facilitate a rapid switch back to proliferation under favorable conditions through coordinated induction of E2F1/2, and potentially other targets, ultimately relieving repression of pro- proliferative E2F gene expression and driving cell-cycle entry ^16^. ZNF768 may thus have evolved in mammals to confer an additional layer of regulation governing the tightly controlled decision between proliferation and cell cycle exit (Fig 8F).

Under conditions of stress that promote cell cycle exit, ZNF768 protein levels are rapidly diminished without corresponding changes in mRNA abundance, indicating that post-translational mechanisms, particularly protein degradation, are a major mode of regulation for ZNF768 ^9^. In line with this, our data demonstrate that in normally cycling cells and in the absence of external perturbation, *ZNF768* transcript levels remain relatively constant throughout the cell cycle. Notably, however, ZNF768 protein is rapidly lost upon entry into mitosis, irrespective of whether it is endogenously expressed or ectopically overexpressed. This suggests that mitotic entry represents a physiological context in which rapid clearance of ZNF768 is imperativeThe observation that ZNF768 mRNA levels remain stable throughout mitosis in an unperturbed cell cycle supports the notion that ZNF768 abundance is regulated primarily at the protein level. The maintenance of a steady source of ZNF768 message in dividing cells may also reflect the need for rapid ZNF768 protein expression after completion of cell division for efficient cell cycle reentry. Collectively, these findings underscore the need for tight temporal control of ZNF768 protein abundance and activity, particularly at the conclusion of each cell cycle, supporting a model in which ZNF768 may play a role in resetting transcriptional programs during cell division (Fig 8F). Whether additional ZNF768 targets beyond E2F1/2 contribute to this process remains to be determined, but given the capacity of ZNF768 to regulate many additional transcriptional targets including transcription factors^7^, we strongly anticipate that this will be the case.

Together, our findings identify ZNF768 as a regulator of cell proliferation, acting as a transcriptional activator controlling both cell cycle entry and progression. Its functions appear to be mediated at least in part through control of pro-proliferative E2F expression, positioning ZNF768 upstream of the major transcriptional hubs of the cell cycle. ZNF768 may thus serve as a critical toggle switch between quiescence and proliferation, with potential implications for both development and tumorigenesis. Additional studies will be required to fully understand how ZNF768 is regulated at mitotic entry, re-expressed at mitotic exit, and integrated into the regulatory network governing cell cycle entry.

## Methods

### Cell Culture and Reagents

The following cell lines were used in this study: hTERT-RPE1, hTERT-RPE1 p53 -/- (a kind gift of Daniel Durocher), HeLa, U87, HCT116, MDA-MB-231, and HEK 293T. All cell lines were obtained from either the American Type Culture Collection (ATCC) or the Coriell Institute and maintained under standard sterile cell culture conditions in complete Dulbecco’s Modified Eagle Medium (DMEM), supplemented with 10% fetal bovine serum (FBS) (Sigma, #F1051) and 1% penicillin-streptomycin (Wisent, #450- 201-EL). The following reagents were used in cell culture experiments: puromycin (1 μg/mL or 10 μg/mL) (Sigma, #P8833), blasticidin S-HCl (2.5 μg/mL) (ThermoFisher Scientific, #A1113903), and doxycycline (100 ng/mL) (Sigma, #D3447).

### Generation of Inducible Stable Cell Lines

Inducible stable cell lines for ZNF768 knockdown were generated via viral transduction. For viral particle production, HEK 293T cells were transfected with the appropriate plasmids. Retroviral particles were produced using gag/pol and CMV VSV- G packaging plasmids, whereas lentiviral particles were produced using psPAX2 and pMD2.G plasmids. Virus-containing supernatants were collected 48 hours post- transfection and filtered through a 0.45 μm filter. Target cells were transduced with the viral supernatants for 24 hours in the presence of 8 μg/mL Polybrene (ThermoFisher Scientific, #TR1003G). Following transduction, cells were refreshed with fresh medium and selected over the following days using either 1 μg/mL or 10 μg/mL puromycin or 2.5 μg/mL blasticidin, depending on the viral construct and cell line used.

### Vectors

Lentiviral shRNAs were obtained from the collection of The RNAi Consortium (TRC) at the Broad Institute. sh_ZNF768_1 (TRCN0000017384), sh_ZNF768_2 (TRCN0000017385) and were cloned in Tet-pLKO-puro (gift from Dmitri Wiederschain, Addgene plasmid #21915). For overexpression of ZNF768 protein, lentiviral constructs were obtained from the collection CCSB Broad Resource. The sequence of this vector can be found at the TRC public website: pLX304_ ZNF768- V5 (ccsbBroad304_12602) and was cloned in pcDNA3-Clover (Addgene plasmid #40259) using pLX304_ ZNF768-V5 as template. Then, the ZNF768-V5 and ZNF768-Clover sequences were subcloned in pCW57-MCS1-P2A-MCS2 (Blast) Tet ON-inducible system (Addgene plasmid #80921).

### Synchronization treatments and protein stability experiments

Synchronization treatments were performed as follows, unless otherwise indicated: Thymidine (2 mM for 16 h; Acros Organics), nocodazole (0.3 uM for 16 h; Sigma- Aldrich), RO-3306 (4 uM for 16h, Sigma-Aldrich), paclitaxel (Taxol, 15 nM for 16 h; Calbiochem). For protein stability experiments, cells were treated with cycloheximide (10 µg/mL) (Santa Cruz Biotechnology, #sc-3508), MG132 (20 nM) (Enzo Life Sciences, #89161-566), actinomycin D (8 µM) (Sigma-Aldrich, #A9415-2MG) for 5 hours prior to lysis.

### Western blotting

All cells were rinsed twice with ice-cold phosphate-buffered saline (PBS) before lysis. Cells were lysed in RIPA lysis buffer (150 mMTris-HCL pH 7.5, 150 mM NaCl, 10 mM NaF, 1% NP-40, and 0.1% Na-deoxycholate) with a protease and phosphatase inhibitor cocktail that included 20 mM β-glycerophosphate, 0.1 mM sodium vanadate, 10 mM sodium pyrophosphate, 1 mg/mL, leupeptin, 1 mg/mL, aprotinin and 1 mM AEBSF. Lysed cells were rotated at 4°C for a minimum of 30 min and followed by centrifugation at 13 000 rpm for 10 min at 4°C. The supernatant was collected and quantified using the BCA assay (Thermo Fisher Scientific) prior to Western blotting. Protein extracts were diluted in SDS-PAGE sample buffer and loaded onto 10% SDS- PAGE gels. Membranes (PVDF, Immobilon-P; MilliporeSigma) were blocked in 5% of milk diluted in PBS and containing 0.05% of Tween-20 for 60 min and then incubated overnight at 4°C with a primary antibody in 5% of milk in PBS. Membranes were washed with PBS containing 0.05% Tween-20 and incubated with the appropriate secondary antibody conjugated to horseradish peroxidase for 1 h at room temperature. After three additional washes, antibody binding was detected with either the Clarity or Clarity Max Western ECL substrate (Bio-Rad) and ChemiDoc MP Imaging System (Bio-Rad).

Tissues were homogenized with Triton X-100 containing lysis buffer (50 mM HEPES, pH 7.4, 2 mM EDTA, 10 mM sodium pyrophosphate, 10 mM sodium β-glycerophosphate, 40 mM NaCl, 50 mM NaF, 2 mM sodium orthovanadate, 1% Triton X- 0, and one tablet of EDTA-free protease inhibitors per 25 ml) supplemented with 0.1% sodium lauryl sulfate and 1% sodium deoxycholate. Lysed tissues were rotated at 4 °C for 10 min and then the soluble fractions of cell lysates were isolated by centrifugation for 10 min in a microcentrifuge. Protein levels were then quantified using Bradford reagent and analyzed by Western blotting. Equalized protein extracts were diluted in sample buffer, denaturated by heat (95 °C) for 10 min and loaded on precast gels (Life Technologies). Proteins were transferred to PVDF membranes blocked in 5% milk diluted in PBS-Tween and incubated with their primary antibody overnight at 4 °C. Membranes were washed with PBS containing 0.05% Tween-20 and incubated with the appropriate secondary antibody conjugated to horseradish peroxidase for 1 h at room temperature. After three additional washes, antibody binding was detected with either the Clarity or Clarity Max Western ECL substrate (Bio-Rad) and ChemiDoc MP Imaging System (Bio-Rad).

The following antibodies were used for western blotting: Cyclin A (BD Transduction Laboratories, #AB_398797); Cyclin B1 (Cell signaling, #12231T); Cyclin D1 (New England Biolabs, #2978S); anti-GAPDH (Novus Biologicals, # NB300-221); p21 (Cell Signalling Technology, Cell #2947); ZNF768 (Aviva Systems Biology, FLJ23436) or a previously described ZNF768 antibody ^7^; phospho-Rb (T826) (Abcam, #ab133446), Rb (New England Biolabs, #9313T); E2F1 (Cell Signaling Technology, #3742S); FOXM1 (Abcam Canada, #AB207298-1001); PLK1 (described previously in *Genes & Development* (*Elowe et al., 2007*^30^); CENP-F (Novus Biologicals, #NB500-101C2); KI- 67 (Millipore, #AB9260) . Peroxidase-AffiniPure Goat Anti-Rabbit and Anti-mouse IgG (H+L) were purchased from Jackson ImmunoResearch Inc (Cat #111-035-003, 115- 035-003). ChemiDoc MP Imager and ChemiDoc and Image Lab software (version 6.0) were used to acquire and analyze images.

### SA-β-gal activity assay

Senescence-associated β-galactosidase (SA-β-gal) assays were performed as previously described^41,42^. Cells were fixed with 0.5% glutaraldehyde in PBS for 15 min, then washed and kept in PBS containing 1 mM of MgCl_2_, for at least 24 h. Staining was performed at 37 °C using a solution containing X-Gal, potassium ferricyanide, potassium ferrocyanide and MgCl_2_ in PBS then cells were washed 3 times with H_2_O. Images were taken and the percentage of SA-β-gal positive cells was quantified.

### FACS

Cells were treated with 0.1 μg/ml of doxycycline for specified knockdown time. Cells were then harvested and fixed using cold 80% ethanol as described before^43^. 2.5 × 10^7^ cells/ml were incubated 30 min with WASH buffer (PBS with 1% FBS, 0.09% NaN_3_ pH7.2) containing a FITC-KI67 antibody (Becton Dickinson, # 556026). Cells were then pelleted by centrifugation and the medium was aspirated. Cells were washed in WASH buffer and the final pellet was resuspended in WASH buffer containing propidium iodide (Becton Dickinson, # 556463) and flow cytometry analysis was followed.

Acquisition and analysis were performed on a BD FACSymphony flow cytometer (Becton Dickinson) equipped with BDFACS Diva Software version 9.0.2 (Becton Dickinson). For fluorescence detection of FITC and PI, a blue laser (488 nm) with a 530/30 bandpass filter and a yellow-green laser (561 nm) with a 610/20 bandpass filter were respectively used. Cell cycle analysis was performed using FCS express (De Novo Software).

### Quantitative real-time PCR

Total mRNA was isolated from tissues using the RNeasy Lipid Tissue Mini Kit (Qiagen, 74104). Total mRNA was isolated from cells using the Quick-RNA Purification Kit, Miniprep, Capped Columns (Zymo Research, #R1055). RNA concentration was estimated from absorbance at 260 nm. cDNA synthesis was performed using the iScript^™^ Advanced cDNA Synthesis Kit for RT-qPCR (Bio-Rad). mRNA extraction and cDNA synthesis were performed following the manufacturer’s instructions. cDNA was diluted in DNase-free water (1:10) before quantification by real-time PCR. mRNA transcript levels were measured in duplicate samples using CFX96 or CFX384 touch real-time PCR (Bio-Rad, Mississauga, ON, Canada). Chemical detection of the PCR products was achieved with SYBR Green (Bio-Rad, 172-5271). At the end of each run, melt curve analyses were performed. Gene expression was corrected for the expression level of reference gene. Normalization was done against *B2M*, unless otherwise stated. The primer sequences used are presented in Supp. table 4.

### RNA-seq

Total RNA was isolated form RPE1 cells using the Quick-RNA Purification Kit, Miniprep, Capped Columns (Zymo Research, #R1055). mRNA sequencing libraries were prepared using the NEBNext Ultra II Directional RNA Library Prep Kit for Illumina (New England Biolabs Inc., Ipswich, MA, USA), following the manufacturer’s instructions. In summary, NEBNext Poly(A) mRNA Magnetic Isolation Module (New England Biolabs Inc.) was first used to isolate poly(A)+ RNA from 1 μg of total RNA which was then used as a template for cDNA synthesis via reverse transcription with random primers. Strand specificity was achieved by substituting dTTP with dUTP during second-strand synthesis. The resulting cDNA was converted into double-stranded DNA and underwent end repair. Adapter ligation was followed by purification using the AxyPrep Mag PCR Clean-up Kit (Axygen, Big Flats, NY, USA). Strands containing dUTP were selectively excised, and the libraries were enriched by 9 cycles of PCR to incorporate indexed adapters for multiplexing. Library quality was assessed using a DNA ScreenTape D1000 on a TapeStation 4200, and quantification was performed with a Qubit 3.0 Fluorometer (ThermoFisher Scientific, Canada). Indexed libraries were then pooled in equimolar ratios and sequenced using paired-end 100 bp reads on an Illumina NovaSeq 6000 flow cell at the Next-Generation Sequencing Platform, Genomics Center, CHU de Québec–Université Laval Research Center, Québec City, Canada. The average coverage per sample was approximately 25 million paired end reads. Raw and processed data is available through the NCBI CEO platform under the accession number GSE337127.

### RNA-seq Data Processing

Raw sequencing reads (FASTQ files) were processed and analyzed using the Galaxy platform^44^. Sequencing quality was assessed prior to downstream analysis using standard quality control metrics. Adapter sequences and low-quality bases were trimmed before alignment. Processed reads were aligned to the human reference genome (build hg38) using STAR^45^ and quantification of the number of reads per gene was assessed using FeatureCounts^46^. Differential gene expression analysis was performed using DESeq2^47^ using three biological replicates per condition. Genes with an adjusted p-value < 0.05 and an absolute log2 fold change > 1 were considered significantly differentially expressed. Pathway enrichment analysis was performed for KEGG and REACTOME pathways using the GSVA R package (v2.0.5)^48,49^.

### mRNA stability assay

Cell lines (control, Sh-ZNF768_1 and Sh-ZNF768_2) were seeded in 6-cm dishes. After 24 hours, induction of transgenes was performed by addition of 100 ng/ml of doxycycline (Sigma, #D3447). After two days, cells were treated with or without Actinomicyn D (Millipore Sigma, A9415) at 5 μg/mL. Cell pellets at 0, 2, 4 and 6 hours were acquired. RNA extracted by the Quick-RNA Purification Kit, Miniprep, Capped Columns (Zymo Research, #R1055) and DNase treated and purified via RNA clean- up columns (Zymo Research, RNA Clean & Concentrator-5 (with DNase), VWR; 76020-604). RNA was then reverse-transcribed for RT-qPCR using iScript™ gDNA Clear cDNA Synthesis Kit (Bio-Rad #1725035) following the manufacturer’s instructions. All RT-qPCR analyses were performed on CFX384 Touch Real-Time PCR Detection system instrument (Bio-Rad) using SYBR Select Master Mix (ThermoFisher Scientific; 4472919). The reaction mix (10 μl) was prepared according to the manufacturer’s instructions, using each primer at a final concentration of 300nM. The cycling conditions were set according to the manufacturer’s instructions, using a primer annealing temperature of 58°C. All RT-qPCR reactions were performed in technical triplicates. Data are normalized to expression of the *B2M* gene unless stated otherwise.

### ChIP-seq processing

Publicly available ZNF768 ChIP-seq raw data from four human cell lines were retrieved: Raji and U2OS (GEO accession GSE111879, samples GSM3043267 and GSM3043268 for Raji, GSM3043270 and GSM3043271 for U2OS)^7^, HEK293 (GEO accession GSE76496, sample GSM2026871)^50^ and HepG2 (ENCODE experiment ENCSR181ABP)^51^. Processing of the raw sequencing reads was performed as described^52^. Briefly, initial read quality was assessed with FastQC (v0.12.1)^53^. Reads were screened for overrepresented sequences with fastp (v0.24.0)^54^. Subsequent read processing and alignment were carried out using the ChIP-seq pipeline from GenPipes (v6.1.1)^55^. Reads were first trimmed with Trimmomatic (v0.39)^56^. High-quality reads were aligned to the human reference genome GRCh38 using bwa-mem2 (v2.2.1)^57^, and PCR duplicates were marked with sambamba (v0.8.1)^58^.

For datasets with two biological replicates (Raji, U2OS and HepG2), replicate alignments were pooled with samtools merge (v1.22.1)^59^ and indexed; the single HEK293 replicate was used directly. Signal track bigWig files were generated with deepTools bamCoverage (v3.5.4)^60^ and normalized to RPKM (reads per kilobase per million mapped reads).

ZNF768 binding site coordinates were retrieved directly from the corresponding public repository. HEK293 peaks, provided only in hg19 coordinates, were converted to GRCh38 reference with the rtracklayer liftOver function^61^ using the UCSC hg19ToHg38 chain file. Genome-browser-style track figures were generated with the plotgardener R package (v1.16.0)^62^.

### Analysis of public ChIP-seq datasets with ChIP ReMap

The binding of transcription factors to the *ZNF768* locus was assessed using ChIP-seq datasets available through the ReMap database^63^. Binding events in the proximal promoter region of *ZNF768* were visualized and analyzed using the ReMap genome browser.

### Immunofluorescence

Cells were grown on coverslips and fixed in PTEMF buffer (0.2% Triton X-100, 20 mM PIPES pH 6.9, 1 mM MgCl_2_, 10 mM EGTA and 4% formaldehyde) for 10 min at room temperature. After fixation, coverslips were blocked with 3% bovine serum albumin (BSA) in PBS-Tween 0.2% for at least 30 min prior to incubation with primary and secondary antibodies for 2 and 1 h, respectively, at room temperature. Antibodies were used at 1 μg/ml, unless otherwise indicated. Antibodies against the following proteins were used: PLK1^30^; Cenp-C (MBL International, #PD030). HOESCHT 33342 (Sigma- Aldrich) was used to stain chromatin. Alexa Fluor-affiniPure series secondary antibodies (Thermo Fisher Scientific) or (Jackson ImmunoResearch) were used for immunofluorescence (1:1,000). For live-cell imaging, cells were washed with PBS and treated with SiR-DNA dye (CY-SC007; Cytoskeleton) at 300 nM for 1 h prior to imaging. Secondary antibodies (donkey anti-rabbit, anti-mouse and anti-human Alexa Fluor AffiniPure-488, 594, and 647) were purchased from Jackson ImmunoResearch (Cat #715-545-150, 709-545-149, 711-545-152, 15-585-150, 711-585-152, 709-585-149, 706-605-148, 715-605-151, 709-605-149).

### Confocal microscopy

An inverted Olympus IX81 microscope equipped with a WaveFX-Borealin-SC Yokagawa spinning disc (Quorum Technologies) and an Orca Flash4.0 camera (Hamamatsu) was used for the acquisition of all images. Images were acquired by Metamorph software (Molecular Devices). Optical sections with identical exposure times for each channel within an experiment were acquired and then projected into a single picture using ImageJ. The system was equipped with a motorized stage (ASI) and incubator with atmospheric CO_2_ warmed at 37°C for live-cell imaging. For time-lapse experiments, images were taken with a 20x objective (Olympus UPLSAPO 20x [NA 0.7]), every 3 min. Higher resolution images with acquired with a 100x (UPLSAPO100XO, NA 1.40) objective.

### Human lung samples

The patients included in this study were diagnosed with LUAD and underwent surgical resection at the Institut universitaire de cardiologie et de pneumologie de Québec - Université Laval (IUCPQ-UL). Lung tumors and adjacent normal lung were collected and stored at the IUCPQ-UL site of the Respiratory Health Network Tissue Bank (http://www.tissuebank.ca/). The Research Ethics Committee of IUCPQ-UL approved this study (#2017-2829, 21441) and all participants provided written and informed consent.

### TCGA analysis

TCGAbiolinks R package was used to download mRNA expression data from The Cancer Genome Atlas^64^. Gene expression for each cancer type was converted to logarithmic scale and normalized with mean zero and variance 1 across each row. Pearson correlation analysis was used to determine the strength and direction of associations between variables. Statistical significance was set at p < 0.05.

## Statistical analysis

Statistical analysis and graph plotting were performed in GraphPad Prism Software V10.5.0 and presented as Superplots where applicable^65^. ANOVA tests were used to determine significance, and the exact test depended on the parameters of the experiment. Statistical tests are described in the figure legends for each experiment.

## Acknowledgements

We thank the CHU de Québec Genomics platform and the Cytometry platform for assistance with experiments. SE holds a Canada Research Chair in Mitotic Signalling and Aneuploidy. ML holds a distinguished scholar award and Philippe Joubert a J2 salary award from the Fonds de recherche du Québec (FRQ). Research in the Elowe lab is funded by grants from the Natural science and engineering Council of Canada, the Canadian Institutes of health research, and the FRQ.

## Author contributions

RD, BB, SR, EIJL, AP, AK, DC, FL, RV, and CG designed and /or executed experiments; LT assisted with figure preparation and statistical analysis; RD prepared the initial draft of the manuscript. SH, PJ, ML and SE were responsible for funding acquisition, study design, supervision and final manuscript preparation.

## Supplemental Figures

**Supplementary figure S1.**
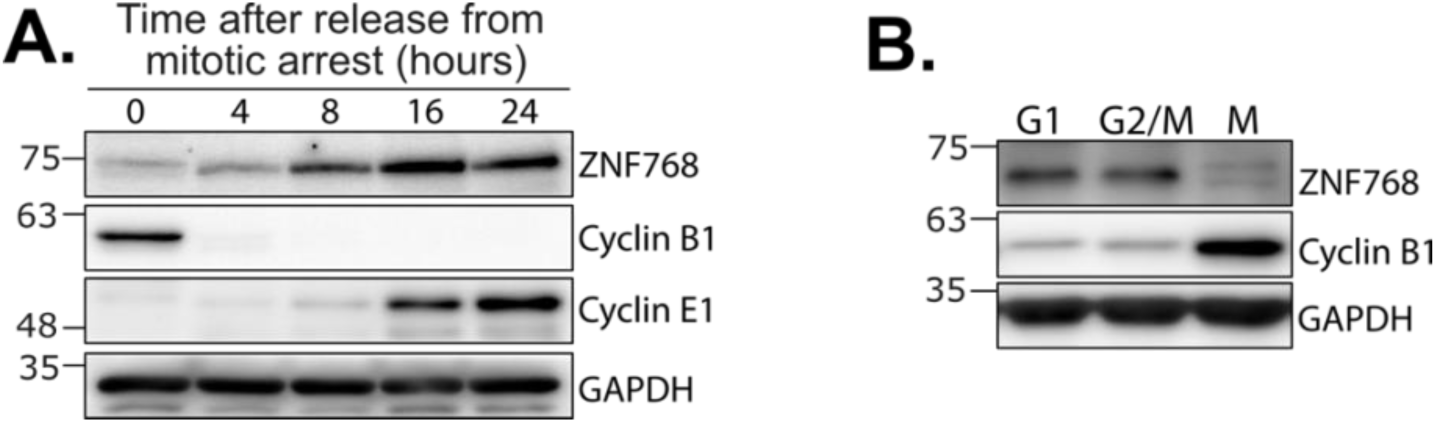
Regulation of ZNF768 protein levels across the cell cycle. **A** Cells were released from a nocodazole-mediated mitotic arrest and lysates were blotted with the indicated antibodies (*n*=3). **B** ZNF768 protein levels in cells synchronized at different stages of the cell cycle: G1 (thymidine); G2/M (RO-3306) and M (nocodazole) (*n*=3).

**Supplementary figure S2.**
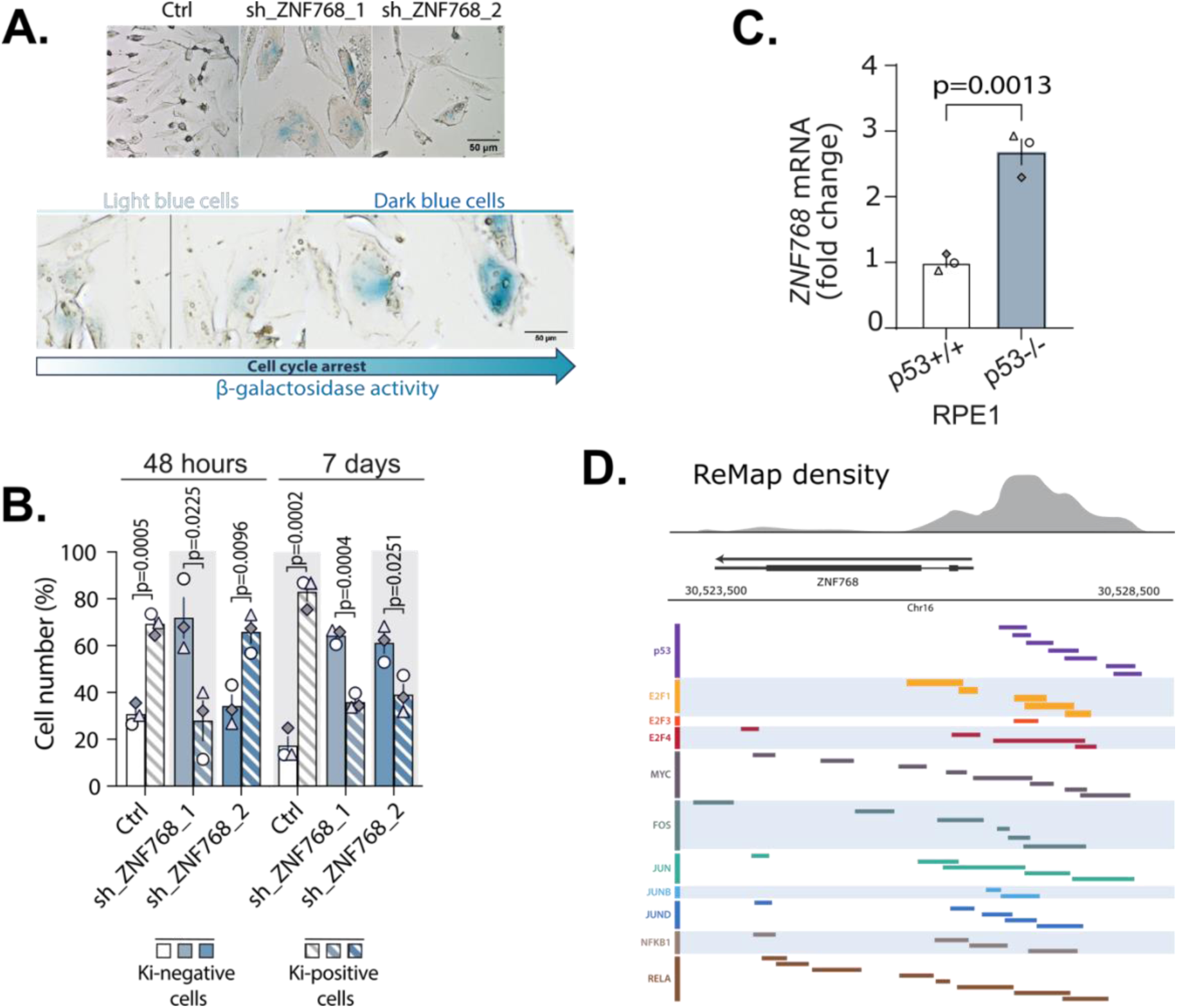
ZNF768 depletion results in cell cycle exit. **A** Representative image of β-galactosidase staining in ZNF768 depleted cells. **B** FACS analysis of PI stained p53+/+ RPE1 cells depleted of ZNF768 for 48 hours or 7 days. **C** Relative expression of *ZNF768* mRNA in p53+/+ and p53-/- RPE1 cells. **D** Analyses of ChIP-Seq experiments showing the binding of different transcription factors close to the promoter of *ZNF768* generated using ReMap. In panel **B** and **C**, significance was determined by unpaired *t* test.

**Supplementary figure S3.**
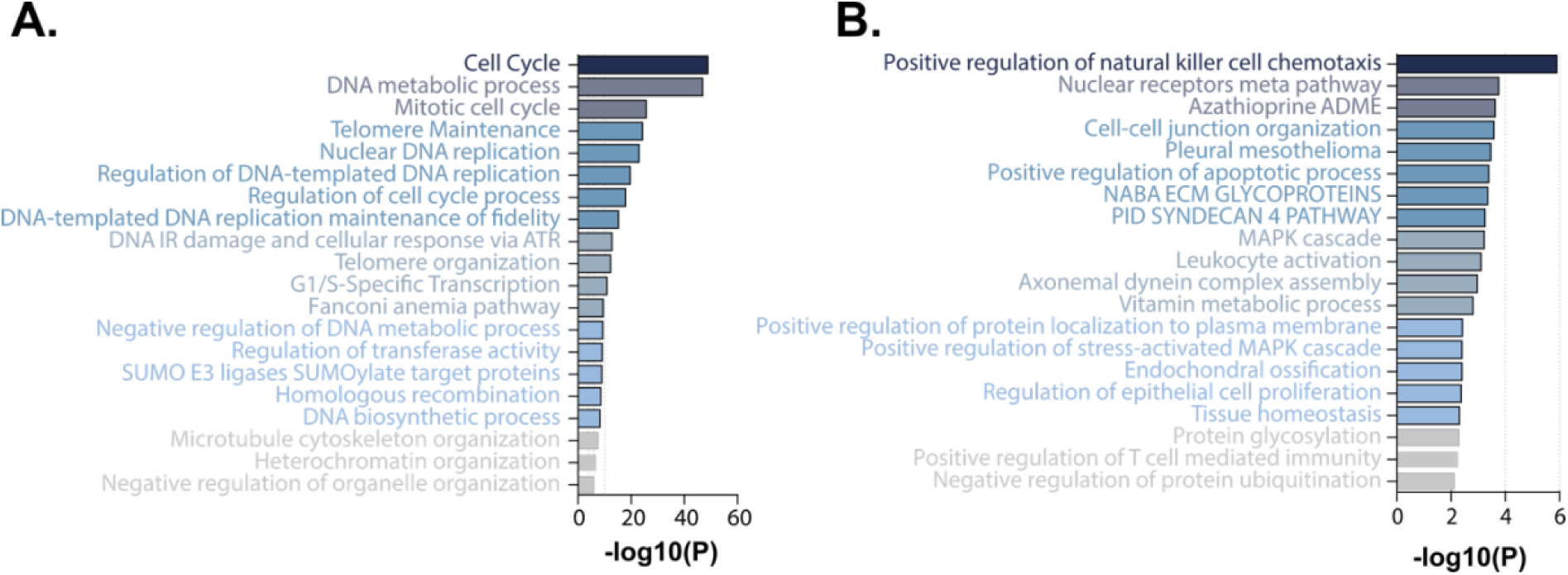
Gene Ontology enrichment of differentially expressed genes inZNF768 depleted cells. **A** Gene ontology analysis performed with Metascape on the genes identified by RNA-seq to be commonly downregulated in response to ZNF768 depletion with sh_ZNF768_1 and sh_ZNF768_2. **B** Gene ontology analysis performed with Metascape on the genes identified by RNA-seq that were commonly upregulated in response to ZNF768 depletion with sh_ZNF768_1 and sh_ZNF768_2.

**Supplementary figure S4.**
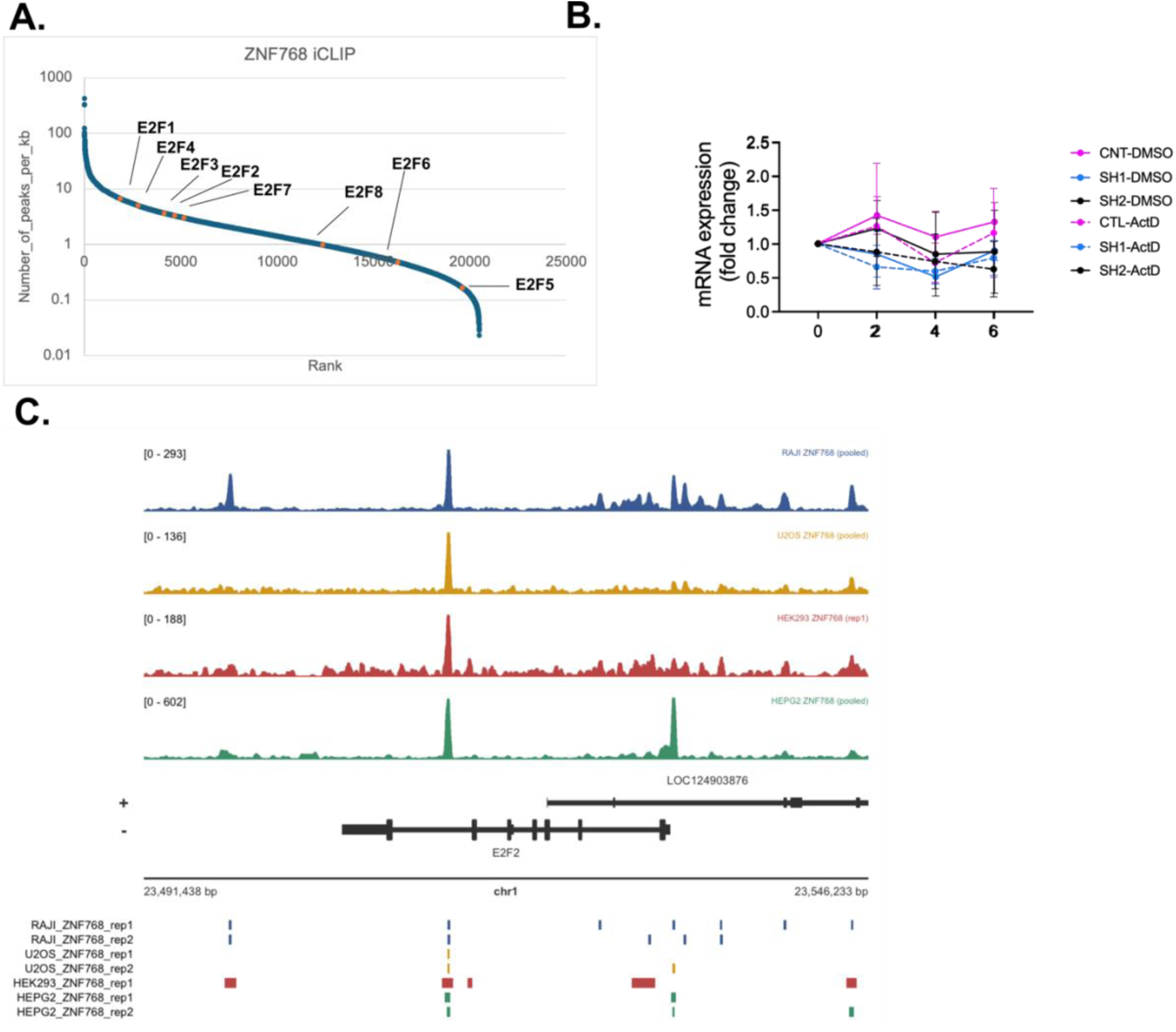
E2F1 is downstream target of ZNF768. **A** iCLIP binding of ZNF768 with mRNA from Nabeel-Shah *et al*. **B** Stability of *E2F1* transcripts in control or ZNF768-depleted RPE1 cells after 2, 4 or 6 hour treatment with cycloheximide (*n*=3). **C** ZNF768 ChIP tracks at the *E2F2* genomic locus.

**Supplementary figure 5.**
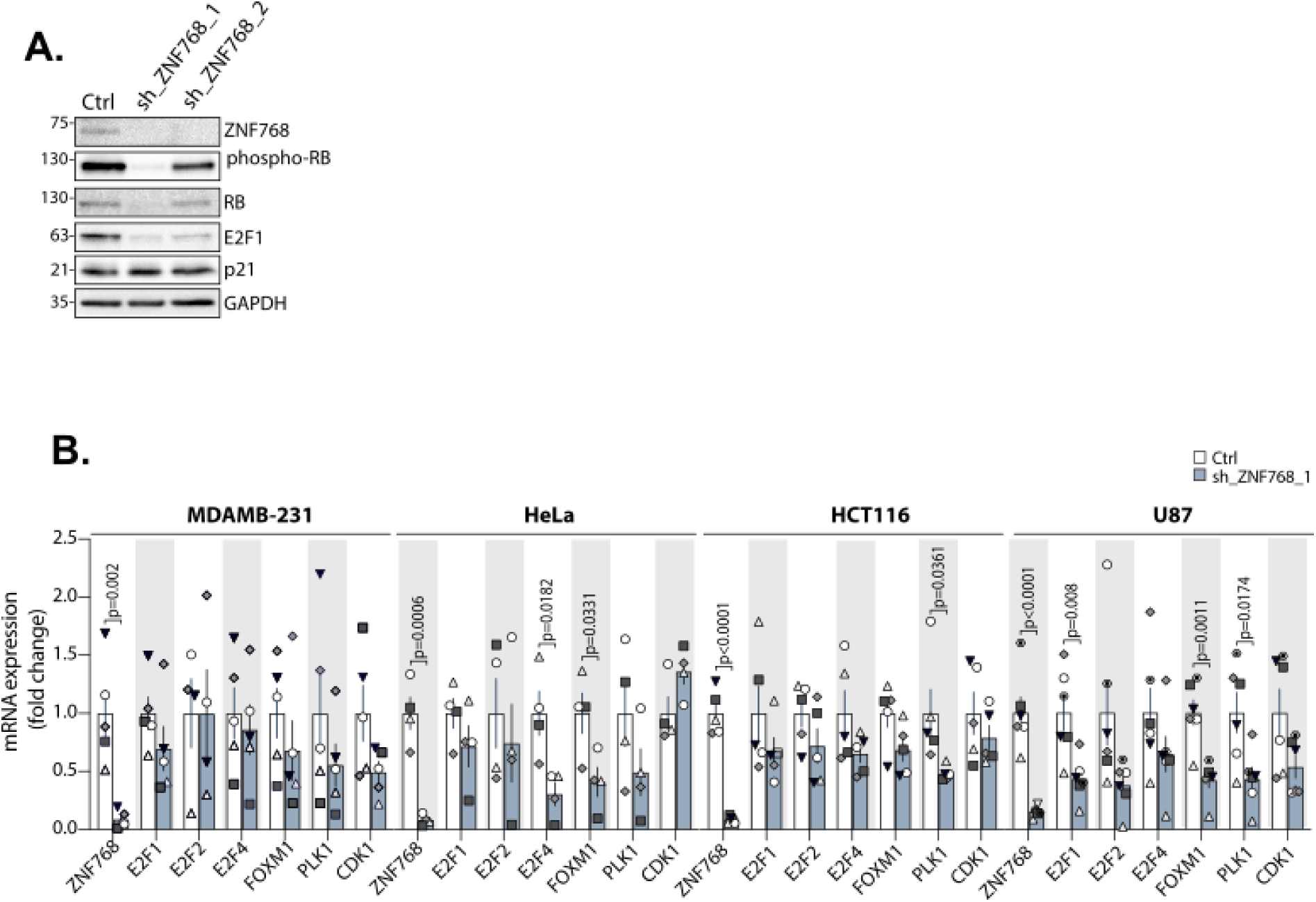
ZNF768 depletion downregulates E2F1 and E2F1target genes in multiple cell lines. **A** Lysates from control and ZNF768 depleted p53+/+ RPE1 cells were blotted for the indicated antibodies (*n*=3). **B** RT-qPCR analysis of the indicated genes after induction of ZNF768 depletion in a panel of cancer cell lines (*n*= 4-6). In panel **B**, significance was determined by unpaired *t* test.

**Supplementary Table S1. Differentially expressed genes identified by DESeq2 following ZNF768 knockdown (adjusted P value < 0.05 and |log2FC| > 1).**

List of genes significantly differentially expressed in response to ZNF768 knockdown with both shZNF768 conditions, as identified by DESeq2. Only genes with an adjusted P value < 0.05 and an absolute log2 fold change (|log2FC|) > 1 were retained. The table includes gene identifiers, log2 fold changes, raw P values, and adjusted P values (padj).

**Supplementary Table S2. Metascape enrichment analysis results of upregulated genes.**

Results of the Metascape enrichment analysis performed on the overlapping genes upregulated following ZNF768 knockdown in both shZNF768 conditions. The table lists the significantly enriched biological processes, molecular functions, cellular components, pathways, and other functional annotations, together with the corresponding enrichment statistics, including gene counts, enrichment factors, P values, and false discovery rate (FDR)-adjusted P values where applicable.

**Supplementary Table S3. Metascape enrichment analysis results of downregulated genes.**

Results of the Metascape enrichment analysis performed on the overlapping genes downregulated following ZNF768 knockdown in both shZNF768 conditions. The table lists the significantly enriched biological processes, molecular functions, cellular components, pathways, and other functional annotations, together with the corresponding enrichment statistics, including gene counts, enrichment factors, P values, and false discovery rate (FDR)-adjusted P values where applicable.

**Supplementary Table S4. Primers used for RT-qPCR.** The table lists the forward and reverse primer sequences used for RT-qPCR analysis in this study.

## Notes

### Competing Interest Statement

The authors have declared no competing interest.

